# Integration of Gene Expression and DNA Methylation Data Across Different Experiments

**DOI:** 10.1101/2022.09.21.508920

**Authors:** Yonatan Itai, Nimrod Rappoport, Ron Shamir

## Abstract

Integrative analysis of multi-omic datasets has proven to be extremely valuable in cancer research and precision medicine. However, obtaining multimodal data from the same samples is often difficult. Integrating multiple datasets of different omics remains a challenge, with only a few available algorithms developed to solve it.

Here, we present INTEND (IntegratioN of Transcriptomic and EpigeNomic Data), a novel algorithm for integrating gene expression and DNA methylation datasets covering disjoint sets of samples. To enable integration, INTEND learns a predictive model between the two omics by training on multi-omic data measured on the same set of samples. In comprehensive testing on eleven TCGA cancer datasets spanning 4329 patients, INTEND achieves significantly superior results compared to four state-of-the-art integration algorithms. We also demonstrate INTEND’s ability to uncover connections between DNA methylation and the regulation of gene expression in the joint analysis of two lung adenocarcinoma single-omic datasets from different sources. INTEND’s data-driven approach makes it a valuable multi-omic data integration tool.

The code for INTEND is available at https://github.com/Shamir-Lab/INTEND.

## 1 Introduction

Emerging technological advances in recent years have made high throughput genome-wide sequencing a central tool for biological research. It allows the collective analysis of various types of biological data (commonly termed ‘omics’), in a single tissue or even at the level of a single cell. These include genomics – covering the DNA sequence itself; transcriptomics – the expression levels of genes in the form of messenger RNAs; epigenomics –reversible modifications on the genetic data, e.g. DNA methylation and chromatin accessibility; proteomics – the levels of translated proteins; and more. Although the analysis of a single omic may generate meaningful insights, it may be necessary to conduct a multi-omic integrative analysis to comprehensively understand a biological system and its complexities. For brevity, will use throughout the term *integration* for integrative analysis. Hence, integrating different omic datasets is one of the most interesting challenges in computational biology today, with the potential of opening new avenues in cancer research and precision medicine (Chakraborty et al. 2018; Efremova and Teichmann 2020; “Method of the Year 2019: Single-Cell Multimodal Omics” 2020)

### 1.1 Multi-omic integration – diverse problems, diverse approaches

One way to obtain multi-omics data for analysis is to simultaneously measure more than one omic from the same tissue. For example, TCGA (The Cancer Genome Atlas) (McLendon et al. 2008) contains multimodal data for numerous tissues spanning dozens of cancer types. The main data types covered by TCGA are genotype, copy number variations, genome methylation, mRNA expression, and miRNA expression, along with clinical data. Multimodal data can be also obtained at the cell level by simultaneously measuring multiple types of molecules within the cell (Angermueller et al. 2016; Clark et al. 2018; Argelaguet et al. 2019). Such technologies are relatively new and expensive, and thus so far there is much less data of multiple omics from the same cells.

Schematically, we can categorize the integration problems into three scenarios (**Figure 1A**):

a. Single omic – multiple datasets (SO/MD). Here only one omic type is used but multiple datasets (typically experiments from different labs or studies) need to be analyzed together.
b. Multiple omic – single dataset (MO/SD). Here there is one set of samples on which several omics were measured, and the feature sets of the different omics are disjoint.
c. Multiple omics – multiple datasets (MO/MD). This problem generalizes both (a) and (b).

**Figure 1.**
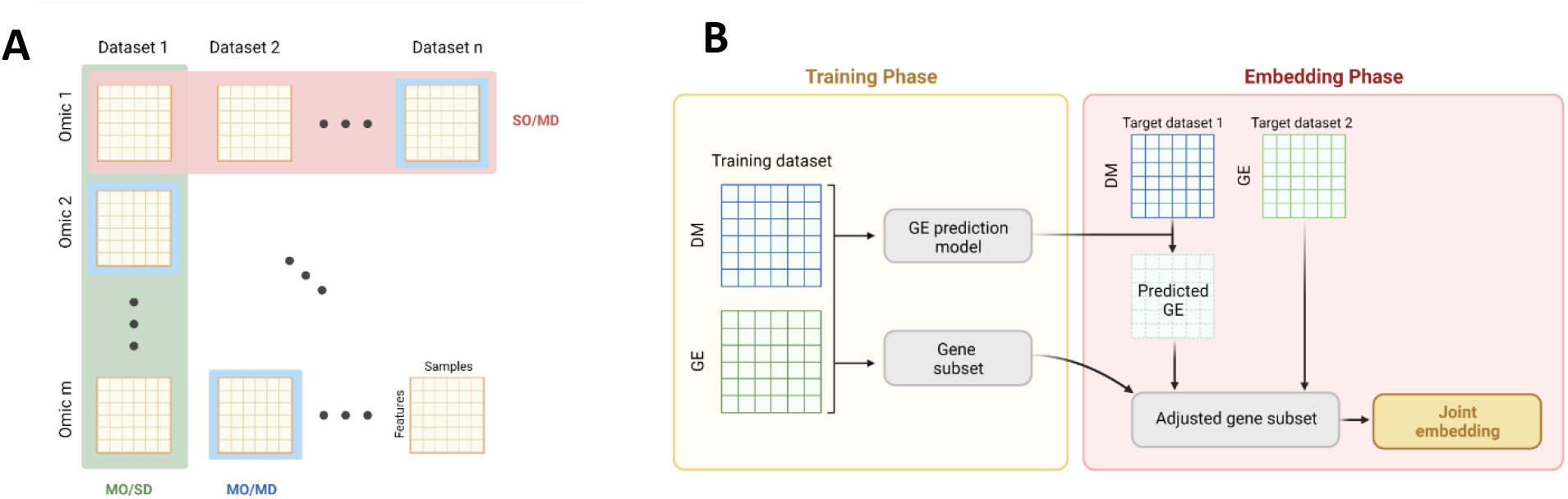
(A) Three scenarios of integration problems: Green: single omic – multiple datasets (SO/MD); red: multiple omic – single dataset (MO/SD); blue: multiple omics – multiple datasets (MO/MD). (B) An overview of the two phases of INTEND: the training phase and the embedding phase.

Many algorithms were developed to handle the integration in the MO/SD setting. These include DIABLO (Singh et al. 2019), iCluster (Shen, Olshen, and Ladanyi 2009), and MOFA/MOFA+ (Argelaguet et al. 2018; 2020), which use latent variable analysis approach; iNMF (Yang and Michailidis 2016), which uses nonnegative matrix factorization; similarity-based methods like SNF (B. Wang et al. 2014), NEMO (Rappoport and Shamir 2019b; 2018) and MONET (Rappoport, Safra, and Shamir 2020); and scAI (Jin, Zhang, and Nie 2020), which specializes in single-cell data. Other algorithms were developed to tackle the integration in the SO/MD setting. These algorithms should balance the tradeoff between the removal of batch effects and the conservation of biological variance (Luecken et al. 2020). Relevant examples are MNN (Haghverdi et al. 2018), Seurat v3 (Stuart et al. 2019), scVI (Lopez et al. 2018), Scanorama (Hie, Bryson, and Berger 2019), LIGER (Welch et al. 2019), Conos (Barkas et al. 2019) and Harmony (Korsunsky et al. 2019).

The challenge we address in this paper is the composition of the two problems discussed above: MO/MD integration. Only a few algorithms have been developed to tackle this challenge. Both LIGER and Seurat v3 were used to integrate different omic datasets of disjoint sets of cells, specifically transcriptome and epigenome datasets. LIGER was shown to integrate scRNA-seq with genome-wide DNA methylation, and Seurat to integrate scRNA-seq with scATAC-seq (measuring chromatin accessibility).

The motivation behind integrating datasets across different experiments arises from the difficulties to obtain multimodal data from the same samples. These difficulties may be technical inabilities, as mentioned in the context of single-cell data, and economical, a significant factor also in the case of bulk sequencing data. An algorithm that can integrate two different omic datasets measured from disjoint sets of samples, could assist researchers in utilizing data that has already been collected in the past, allowing a multi-omic systemic view on the investigated subject. This could increase efficiency, both in time and in cost. Consider the situation where the methylation patterns inside tumors of a specific cancer subtype are being investigated. The multi-omics approach could suggest further inquiry of the epigenome-transcriptome connections, i.e. obtaining mRNA sequencing from every tumor and conducting an integrative analysis of the methylation and gene expression patterns together. As RNA-seq data is widely available for many cancer subtypes, it may be the case that such RNA-seq data is already available for other samples of that cancer subtype. With an algorithm that can integrate RNA-seq and DNA methylation datasets measured on disjoint samples, the researcher could conduct an integrative multi-omic analysis while measuring only the methylation patterns, thus requiring fewer resources.

The algorithms for MO/MD integration can be classified according to the correspondence information they require as input. Some methods require partial correspondence between the samples (either tissues or cells). One example is the semi-supervised correspondence approach of the MAGAN algorithm (Amodio and Krishnaswamy 2018). This approach uses matching pairs of samples from both datasets to learn the correct alignment of the datasets. Other methods, as LIGER and Seurat, require correspondence information between the features of the different omics. Finally, some methods do not require any correspondence information and assume a common underlying structure that is maintained across technologies and omics. Such methods usually belong to the class of machine learning algorithms that solve the unsupervised manifold alignment problem. One algorithm that uses such techniques to integrate single-cell multi-omics data is the maximum mean discrepancy-manifold alignment (MMD-MA) algorithm (Liu et al. 2019) Another algorithm that can jointly embed two datasets, without any correspondence information between their features or samples, is the joint Laplacian manifold alignment algorithm (JLMA) (C. Wang and Mahadevan 2008). Using a method that does not require any correspondence information may sound appealing, but may not perform adequately when the assumed common underlying structure is weak.

In our study, we developed a method for the integration of transcriptomic and epigenomic data across different experiments. We focused on the integration of gene expression and DNA methylation. Specializing in two particular creates a less general method, but allows us to develop a stronger model: we can incorporate the known biological connections between gene expression and DNA methylation.

### 1.2 Associations between DNA methylation and gene expression

The regulation of gene expression allows cells to increase or decrease the production of proteins or RNA. Such adjustments enable response to external changes in the environment and to internal signals within cells. In complex multicellular organisms, the regulation of genes in particular cellular contexts enables the differentiation and proliferation of cells. Epigenetic modifications mainly include DNA methylation and histone protein modifications, which alter the chromatin structure. These modifications are known to be key factors in the regulation of gene expression. In the last two decades, a strong connection has been established between epigenetic modifications and the development of cancer. Hence, the integration of transcriptomic and epigenomic data has the potential to broaden our understanding of the molecular mechanisms orchestrating the regulation of genes, in both normal and malignant tissues.

DNA methylation in mammals occurs almost exclusively in the 5’ position of a Cytosine followed by a Guanine, commonly termed a CpG site. CpG dinucleotides tend to cluster in CpG islands (CGIs), regions with a high frequency of CpG sites. The majority of CpG dinucleotides (75%) throughout the mammalian genomes are methylated (Tost 2010), except for CGIs, which are mostly unmethylated. About 70% of the proximal promoters of human genes contain a CGI, and reciprocally, about 50% of the CGIs are located near a gene’s transcription start site (TSS). In fact, CGIs are strongly linked to the regulation of transcription (Deaton and Bird 2011). Although CGIs are mostly hypomethylated, there are known examples of their methylation, resulting in stable silencing of the associated promoter. However, it is believed that CGI methylation does not initiate the silencing of genes, but assists in making the silenced state permanent (Deaton and Bird 2011). For example, in X chromosome inactivation, the methylation process of CGIs in the X chromosome has been shown to start only after gene silencing. However, when DNA methylation is inhibited, genes in the X chromosome can be reactivated.

The connection between CGI hypermethylation and silencing of genes is not the only relationship observed between methylation and gene expression. There is evidence of both strong positive and strong negative correlations between gene-body methylation and gene expression (Jjingo et al. 2012). Other studies have shown that hypermethylation of CGIs in cancer tissues is not always accompanied by a decrease in gene expression (Moarii et al. 2015). These findings suggest that DNA methylation can play diverse roles in gene regulation, depending on the genomic context (Bhasin et al. 2015). This should be considered when using multi-omic integration algorithms like LIGER and Seurat, which require correspondence information between the features of the different omics. The methods that are currently used to link the feature spaces of DNA methylation and gene expression assume a simplistic connection between the two (see LIGER description in the Supplement). The complex and not fully understood relationship between DNA methylation and gene expression stresses the necessity for a more sophisticated approach.

### 1.3 Our approach

In this paper, we present a novel algorithm for the MO/MD problem. The algorithm is called INTEND (IntegratioN of Transcriptomic and EpigeNomic Data). Specifically, INTEND aims to integrate gene expression (GE) and DNA methylation (DM) datasets covering *disjoint* sets of samples. INTEND does not use any correspondence information between the samples in the two datasets (e.g. knowing which GE and DM profiles originated from the same individual). To handle the complex connections between DM and GE, INTEND learns a predictive model between the two, by training on multi-omic data measured on the same set of samples. To the best of our knowledge, this is the first use of a predictive model in the context of the studied problem.

As a preliminary step, for each gene, INTEND learns a function that predicts its expression based on the methylation levels in sites located proximal to it. To integrate the target methylation and gene expression datasets, INTEND first predicts for each methylation profile its expression profile. Then, it identifies a set of genes that will be used for the joint embedding of the expression and predicted expression datasets. At this stage, both datasets share the same feature space. INTEND then employs canonical-correlation analysis (CCA) to jointly reduce their dimension.

We evaluated the performance of INTEND by comparing it to four state-of-the-art MO/MD integration methods: LIGER, Seurat v3, JLMA, and MMD-MA. The first two require correspondence information between the different omic features, in order to create a common feature space before the integration, whereas the last two do not require such information. We used eleven TCGA cancer datasets spanning 4329 patients for testing the algorithms in multiple integration tasks. We also showed the utility of the method in identifying SKCM cancer subtypes and in joint analysis of LUAD using two single-omic datasets obtained from different individuals.

## 2 Materials and Methods

### 2.1 INTEND algorithm

INTEND works in two phases (**Figure 1B**). The training phase receives as input training data consisting of GE and DM profiles measured on the same set of samples. The algorithm uses this data to learn the connections between the omics. This will allow it later to make accurate predictions of expression levels of specified genes based on a given methylation profile. The training process can be executed once for any number of future integration tasks. Intuitively, the multimodal data used in the training process should be “biologically similar” to the datasets that INTEND will integrate subsequently. However, as we shall show, even when we used INTEND to integrate datasets covering tumor types that were different from the ones covered by the multimodal training data, it performed well.

For the embedding phase, INTEND’s inputs are from two disjoint cohorts, denoted T1 and T2. They include a DM matrix for T1 and a GE matrix for T2. It proceeds in three steps: (1) Creation of predicted GE matrix for T1 based on the DM data. (2) Selection of a subset of the genes based on the predicted GE for T1, the GE for T2, and the trained model from the preliminary step. (3) Reducing jointly the dimension of the two GE datasets on the selected gene set.

#### 2.1.1 The training phase

The preliminary training phase aims to learn connections between GE and DM using training data. Its inputs are expression and methylation profiles for the same set of *n* samples. *E_train_* is an |*f_E_*| × *n* expression matrix, where *f_E_* is the set of genes for which the expression was measured. The methylation matrix *M_train_* has dimensions |*f_M_*| × *n*, where *f_M_* is the set of measured methylation sites. The goal is to determine a function for every gene *g*, that predicts the expression level of *g* based on the methylation levels of potentially relevant sites. Let 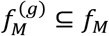 be the set of relevant sites (its creation is described below). For a methylation profile 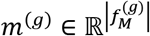, we seek a function 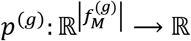, s.t. *p*^(*g*)^ (*m*^(*g*)^) is the predicted expression level of *g*.

##### Model

We hypothesized that accurately predicting the expression levels of even a small number of genes, from an input methylation matrix, will enable successful integration. To achieve this goal, we developed a prediction model considering the known connections between methylation in promoter CGIs and gene expression (Deaton and Bird 2011), as well as gene-body methylation (Jjingo et al. 2012). Furthermore, to capture the variation in the correlation between methylation and expression across the CGI, its shores and shelves, and also outside CGIs (Moarii et al. 2015), the model uses the methylation levels in each probe separately.

For each *g* ∈ *f_E_* we set 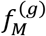 to be all the probed methylation sites in the range 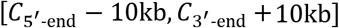, where *C*_5’-end_ and *C*_3’-end_ are the coordinates of *g*’s 5’-end and 3’-end on the chromosome, respectively. While in certain cases more distal methylation sites were reported to affect gene expression (Aran, Sabato, and Hellman 2013), the main effect is usually due to proximal sites (Deaton and Bird 2011). We limited the range in order to have modest-size gene models. As we will show, such models provide a good basis for the integration task.

The size of 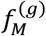 may vary due to the variability in gene length and the assay’s coverage. Genes that had less than two measured methylation sites were removed from the model. For example, in a TCGA training set that we used, spanning 10 cancer subtypes (the datasets listed in **Table 1**, excluding LUAD) and spanning 3852 tumor samples, 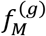 contained 25 sites on average, a median of 19, and a maximum of 1055 sites (**Supplementary Figure 1**). Let 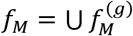 the union of the used methylation sites for all genes.

**Table 1.** Summary information of TCGA cancer datasets used.

| Cancer type | Abbreviation | Number of patient samples |  |  |
| --- | --- | --- | --- | --- |
|  |  | Gene expression | DNA methylation | Both |
| Acute Myeloid Leukemia | **AML** | 173 | 194 | **170** |
| Bladder Urothelial Carcinoma | **BLCA** | 427 | 440 | **425** |
| Colon Adenocarcinoma | **COAD** | 328 | 353 | **298** |
| Brain Lower-Grade Glioma | **LGG** | 534 | 534 | **530** |
| Liver Hepatocellular Carcinoma | **LIHC** | 424 | 430 | **414** |
| Lung Adenocarcinoma | **LUAD** | 576 | 507 | **477** |
| Pancreatic Adenocarcinoma | **PAAD** | 183 | 195 | **183** |
| Prostate Adenocarcinoma | **PRAD** | 550 | 553 | **533** |
| Sarcoma | **SARC** | 265 | 269 | **263** |
| Skin Cutaneous Melanoma | **SKCM** | 473 | 475 | **473** |
| Thyroid Carcinoma | **THCA** | 572 | 571 | **563** |

For each *g*, after obtaining 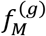, INTEND uses Lasso regression model (Tibshirani 1996; Friedman, Hastie, and Tibshirani 2010) to learn the prediction function *p*^(*g*)^ and select model features. Lasso was run using the glmnet R package and the optimal value of the penalty constant was chosen using 10-fold crossvalidation on the training set. Using Lasso allows the preliminary step to handle genes with a large number of methylation sites, by ignoring sites that have little relevance for the gene expression prediction. For example, in training on the 10 cancer subtypes mentioned above, the maximal number of probes with non-zero coefficients per gene was 424, and the median was 12 (**Supplementary Figure 2**).

After calculating *p*^(*g*)^ for every *g* in every training sample, the 2000 genes with the highest *R*^2^ between predicted and observed gene expression are identified for use in the next stages of INTEND. For example, using the above training set, the average *R*^2^ of all 19143 genes considered was 0.30, and the average *R*^2^ of the top 2000 genes was 0.68 (**Supplementary Figure 3**).

Note that when applying the preliminary step to certain cancer subtypes, the subsequent algorithmic steps use only data from other subtypes, in order to avoid overfitting.

#### 2.1.2 The embedding phase

The inputs for the main phase of the algorithm are:

1. A DM matrix *M*, for one target set of samples (T1), of dimensions |*f_M_*| × *n_M_*
2. A GE matrix *E* for a second, disjoint target set of samples (T2), of dimension |*f_E_*| × *n_E_*
3. A desired dimension *d* for the shared space

Additionally, the prediction functions *p*^(*g*)^ for each *g* from the preliminary step are used. The requested output is a *d* × (*n_M_* + *n_E_*) matrix denoted *S*, which contains the projections of the input and predicted expression profiles into the shared d-dimensional space. The phase has three steps:

##### Step 1: Gene expression prediction using methylation data

Let *p*^(*g*)^ be the learned prediction function for gene *g* and let *m*_1_,*m*_2_,…*m_n_M__* be the methylation profiles in *M*. Recall that 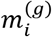 describes the methylation levels of *m_i_* in 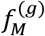 (possibly with some coefficients zeroed by the Lasso process). We apply *p*^(*g*)^ on 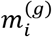 and get the predicted expression 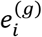. We denote the predicted expression profile for *m_i_* as 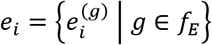. This step results in the predicted expression matrix *P* = (*e*_1_,*e*_2_,…,*e_n_M__*).

##### Step 2: Selecting genes

Denote the 2000 genes selected in the training phase by *G_R_*. The expression of these genes has the highest likelihood to be predicted accurately by the methylation profile, at least in the tissue types and states included in the training set. However, the target datasets may originate from a different tissue type or state. Hence, an additional heuristic for feature selection is employed. Genes may be regulated by mechanisms other than DNA methylation. Thus we assumed that the genes that are most likely to be regulated by the methylation profile are the ones with high variability in both methylation and expression levels. Let *G_E_* denote the 2000 genes with the highest expression variability in *E*. Let *G_P_* denote the 2000 genes with the highest variance in the predicted expression *P*. We select the following genes from *E* and *P*:

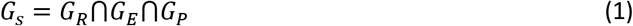

The resulting matrices are *E_G_s__* and *P_G_s’__*, with dimensions |*G_s_*| ×*n_E_* and |*G_s_* | ×*n_M_* respectively. The size of *G_s_* varies depending on the training and target datasets. Finally, each row of *E_G_s__* and *P_G_s__* is centered and scaled separately so that each feature has zero mean expression level and unit variance.

##### Step 3: Embedding

The last step applies CCA to *E_G_s__* and *P_G_s’__*, and produces the integrated matrix *S*. CCA is a dimension reduction method that finds linear combinations of features across datasets such that these combinations have maximum correlation (Hotelling 1936). It was used in the context of computational genomics to project datasets that share the same samples but have different features (the MO/SD setting) to a common lowdimensional feature space. CCA has been used in this way for example in multi-omic clustering (Witten and Tibshirani 2009; Rappoport and Shamir 2018). In contrast, here we apply CCA to *E_G_s__* and *P_G_s__*, which cover samples from different datasets but share the same set of genes *G_s_* (similar to the SO/MD setting). This approach for utilizing CCA was introduced in Seurat v2 (Butler et al. 2018).

Let us denote 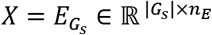 and 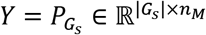. Let *d* ≤ min (*n_E_,n_M_*). CCA aims to find canonical correlation vectors *u*_1_, … *u_d_,v*_1_, …,*v_d_* such that the correlations between the projections *Xu_i_* and *Yv_i_* are maximized, under the constraint that *Xu_i_* is uncorrelated with *Xu_j_* for *j* < *i* and the same for *Yv_i_* and *Yv_j_*. To get the first pair of canonical correlation vectors, the following optimization problem should be solved:

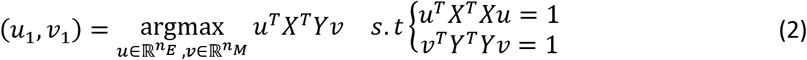

When |*G_s_*| is smaller than the number of samples *n_E_* and/or *n_M_*, the solution for *u*_1_, *v*_1_ is not unique. To overcome this, as proposed in Butler et al., the covariance matrix within each dataset is treated as if it were diagonal, resulting in the following problem:

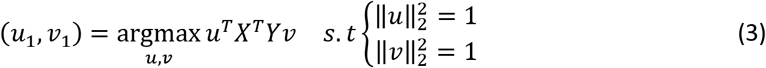

We scale and center the columns of *X* and *Y* to have a mean of 0 and variance of 1 (in the previous step the same process was applied to the rows). The problem can solved using Lagrange multipliers. See the Supplement for details.

The code for INTEND is available at https://github.com/Shamir-Lab/INTEND.

### 2.2 Data

#### 2.2.1 TCGA data

To assess performance, we used RNA-seq and DM data from TCGA (McLendon et al. 2008) covering 11 different cancer types. See **Table 1** for cancer types, their abbreviations and statistics. The data was downloaded using the TCGA-Assembler software (Wei et al. 2018; Zhu, Qiu, and Ji 2014). We used only 4329 samples for which both omics were measured.

The DM data we used was gathered with Illumina’s Infinium HumanMethylation450 BeadChip assay. The levels of > 450,000 methylation sites were reported as *β*-values. The RNA-seq data was gathered with Illumina HiSeq assay, and quantified using RSEM (Li and Dewey 2011). In each GE and DM sample the zero counts were removed, then the raw count values were divided by the 75^th^ percentile of the counts, and then multiplied by 1000. In both omics, we downloaded the data after these transformations from the TCGA website.

#### 2.2.2 An additional LUAD gene expression dataset

In addition to the TCGA LUAD data, we used RNA-seq profiles from 172 tumors of LUAD patients from Singapore (Chen et al. 2020). GE was quantified with RSEM and normalized as done for the TCGA data.

#### 2.2.3 Data preprocessing

To handle missing values, for each dataset, features with > 5% missing values were removed, and then samples with > 5% missing values were removed. Subsequently, the missing values per each feature were imputed to the mean of this feature across all samples. The number of features and samples in each dataset we used, before and after the handling of missing values, are described in **Supplementary Table 1.** Finally, for GE data from all sources and for all purposes, we added 1 pseudo-count to each value and log-transformed the result.

#### 2.2.4 Running other algorithms

We evaluated the performance of INTEND by comparing it to four state-of-the-art MO/MD integration methods: LIGER, Seurat v3, JLMA, and MMD-MA. The methods are briefly described in the Supplement. To use LIGER and Seurat, we supplied the algorithms with an aggregated gene-level methylation matrix as input, as they require correspondence information between features across omics. The aggregated matrix computation process is described in the Supplement. JLMA and MMD-MA algorithms do not require correspondence information between the features. However, empirical results from (Liu et al. 2019) showed that JLMA failed to integrate GE and DM using the local geometry metric as a measure for cross-omic similarity. Hence, we computed the cross-omic similarity matrix for JLMA based on the aggregated gene-level methylation matrix. For MMD-MA we used both the original methylation data and gene-level methylation matrix as inputs. We denoted the runs of JLMA and MMD-MA with the gene-level methylation matrix as JLMA WFCI (with features correspondence information) and MMD-MA WFCI. We ran all the algorithms with their default recommended hyper-parameters, and whenever applicable, we used the algorithm’s pipeline for feature selection and normalization. Since MMD-MA and JLMA do not include a method for feature selection, when running them in the WFCI mode, we selected the *n* genes with the highest variance in expression, for *n* = 500 and 2000. Further details regarding how each of the algorithms was applied, including hyper-parameters and additional necessary preprocessing steps, are described in the Supplement.

### 2.3 Evaluating the quality of the results

For the TCGA data, we have the true pairing of samples that represent different omic measurements of the same patient. This pairing is not given as input to the integration algorithms and can therefore be used to evaluate their results. We use the metric defined in Liu et al. to evaluate the algorithms. For GE and DM input datasets covering *n_E_* and *n_M_* samples respectively, each algorithm produces a *d*-long vector of the projected expression *e_i_* for each sample *i* and a *d*-long vector of the projected estimated expression *m_j_* based on the methylation for each sample *j*. For patient *f_i_*, let be the fraction of samples *j* with projections *m_j_* closer to *e_i_* than *m_i_*. We call it the “fraction of samples closer than the true match” (FOSCTTM). FOSCTTM ranges from 0 to 1, where 0 means that the true match of a sample *i* is the closest to *i* in the projected space. We calculate the FOSCTTM for every sample in the GE and DM datasets, and average these values. A perfect integration will have a score of 0. For a random projection, the expected FOSCTTM is 0.5.

### 2.4 Clustering

For clustering (subsection 3.3), we used the k-means algorithm of Hartigan and Wong (1979), with maximum number of 100 iterations and 100 different starting solutions. We selected the desired number of clusters using the “elbow method” as described in Rappoport and Shamir (2018). Let *v*(*i*) be the total within-cluster sum of squares for a solution with *i* clusters, then we chose *i* for which the point *v*(*i*) had the maximum curvature. Specifically, we chose the *i* that maximized the following approximation of the second derivative of *v*:

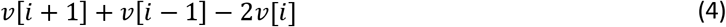

## 3 Results

We applied INTEND in several settings. In the first part, we applied INTEND and four other algorithms in several integration tasks of GE and DM data, using eleven cancer datasets from TCGA. We also demonstrated the utility of the method in identifying SKCM cancer subtypes. In the second part, we used INTEND for the integration of datasets from two different sources, covering two populations of LUAD patients.

Our first set of analyses compared five algorithms: INTEND, LIGER, Seurat v3 (hereafter: Seurat), MMD-MA, and JLMA. We used eleven datasets of different cancer types from TCGA. First, we integrated GE and DM data of the same cancer type, for each of the eleven types. Next, we integrated data of four cancer types simultaneously.

### 3.1 Single cancer dataset integration task

We first ran the algorithms with input datasets of a single cancer subtype. We used the eleven datasets listed in **Table 1**. For each dataset, we considered only the subset of samples measured in both omics. The total number of samples used in these integration tasks was 4329, where dataset sizes ranged from 170 to 563. For each cancer dataset, we trained a new regression model in INTEND’s preliminary phase, using the samples of the remaining ten cancer datasets as the training set. To evaluate the results, we used the pairing information between samples from the two omics measured on the same tissue to calculate the FOSCTTM score.

We ran the algorithms using projected space dimension *d* ranging from 2 to 40, and recorded the best integration scores (average FOSCTTM). The results are summarized in

**Table 2** and **Supplementary Figure 5**. INTEND performed best across all datasets and all *d* values, and substantially better than the rest, with MMD-MA the second performer. In fact, INTEND results were often 1-2 orders of magnitude better than those of all the other methods.

**Table 2.** Average FOSCTTM of algorithms for integrating GE and DM data.

| Cancer/Alg | INTEND | LIGER | Seurat v3 | MMD-MA | MMD-MA<br>WFCI<br>(500) | MMD-MA<br>WFCI<br>(2000) | JLMA<br>WFCI<br>(500) | JLMA<br>WFCI<br>(2000) |
| --- | --- | --- | --- | --- | --- | --- | --- | --- |
| AML | 2.42 (25) | 29.83 (7) | 17.05 (36) | 23.63 (40) | 19.08 (40) | 22.35 (40) | 24.01 (8) | 28.38 (7) |
| BLCA | 0.04 (39) | 39.62 (9) | 13.86 (40) | 11.20 (40) | 16.34 (40) | 14.58 (40) | 34.80 (40) | 37.11 (40) |
| COAD | 0.02 (37) | 26.84 (19) | 19.14 (40) | 12.59 (40) | 12.19 (40) | 12.92 (40) | 32.98 (5) | 34.73 (4) |
| LGG | 6.82 (22) | 41.97 (8) | 32.06 (26) | 8.88 (40) | 15.50 (40) | 12.08 (40) | 37.41 (14) | 32.38 (12) |
| LIHC | 0.14 (36) | 42.34 (3) | 19.23 (38) | 16.04 (30) | 11.02 (30) | 12.94 (30) | 32.68 (21) | 36.03 (12) |
| LUAD | 0.06 (32) | 36.72 (4) | 16.36 (39) | 8.71 (40) | 14.11 (40) | 13.89 (40) | 29.60 (9) | 32.16 (8) |
| PAAD | 0.55 (30) | 36.68 (15) | 24.18 (35) | 11.07 (40) | 23.42 (40) | 16.27 (40) | 29.83 (3) | 27.44 (2) |
| PRAD | 0.37 (38) | 35.96 (8) | 16.32 (17) | 10.88 (40) | 11.15 (40) | 10.99 (40) | 27.14 (2) | 29.53 (2) |
| SARC | 0.05 (35) | 42.06 (15) | 12.86 (36) | 8.86 (40) | 20.97 (40) | 17.42 (40) | 34.47 (7) | 34.73 (5) |
| SKCM | 0.03 (39) | 42.20 (17) | 18.97 (37) | 16.02 (40) | 20.53 (40) | 16.62 (40) | 32.11 (15) | 34.71 (3) |
| THCA | 3.07 (11) | 32.58 (7) | 15.96 (36) | 6.71 (40) | 7.78 (40) | 6.65 (40) | 30.95 (2) | 27.52 (5) |
| Average<br>(all datasets) | 1.23 (31) | 36.98 (10) | 18.73 (34) | 12.24 (39) | 15.64 (39) | 14.25 (39) | 31.45 (11) | 32.25 (9) |

Average FOSCTTM score (percent) for each algorithm on each of the eleven cancer datasets. The optimal score is 0%, and the expected score for a random projection is 50%. The requested shared space dimension $d$ ranges from 2 to 40 for each algorithm. The score shown is the best across all values of $d$ , and the optimal $d$ is written in parenthesis. The numbers 500 and 2000 for MMD-MA and JLMA denote the number of selected genes in the WFCI runs.

In later analyses, we preferred to use the same space dimension *d* for all algorithms. MMD-MA and JLMA do not recommend a method for determining d. For Seurat, the authors originally suggested approaches to select *d* (Butler et al. 2018) but later noted that the identification of this value remains a challenge (Stuart et al. 2019). After running all methods for *d* ∈ [2,40] for all datasets, we observed that most algorithms reach a plateau in the FOSCTTM score around *d* = 40 (**Supplementary Figure 5**). Hence, in subsequent runs we set *d* = 40 for all algorithms, with one exception: LIGER failed to run on the AML dataset with *d* = 40 or *d* = 39, so in that case we used *d* = 38.

Next, we analyzed the FOSCTTM per sample across all methods and datasets. **Figure 2** shows boxplots of the FOSCTTM per sample for each algorithm and cancer dataset using ***d*** = **40**. INTEND’s advantage was prominent, with the entire FOSCTTM interquartile range (***IQR***) at zero for eight of the 11 datasets tested. In six of the 11 datasets, the FOSCTTM was perfect (zero) for > **90**% of the samples.

**Figure 2.**
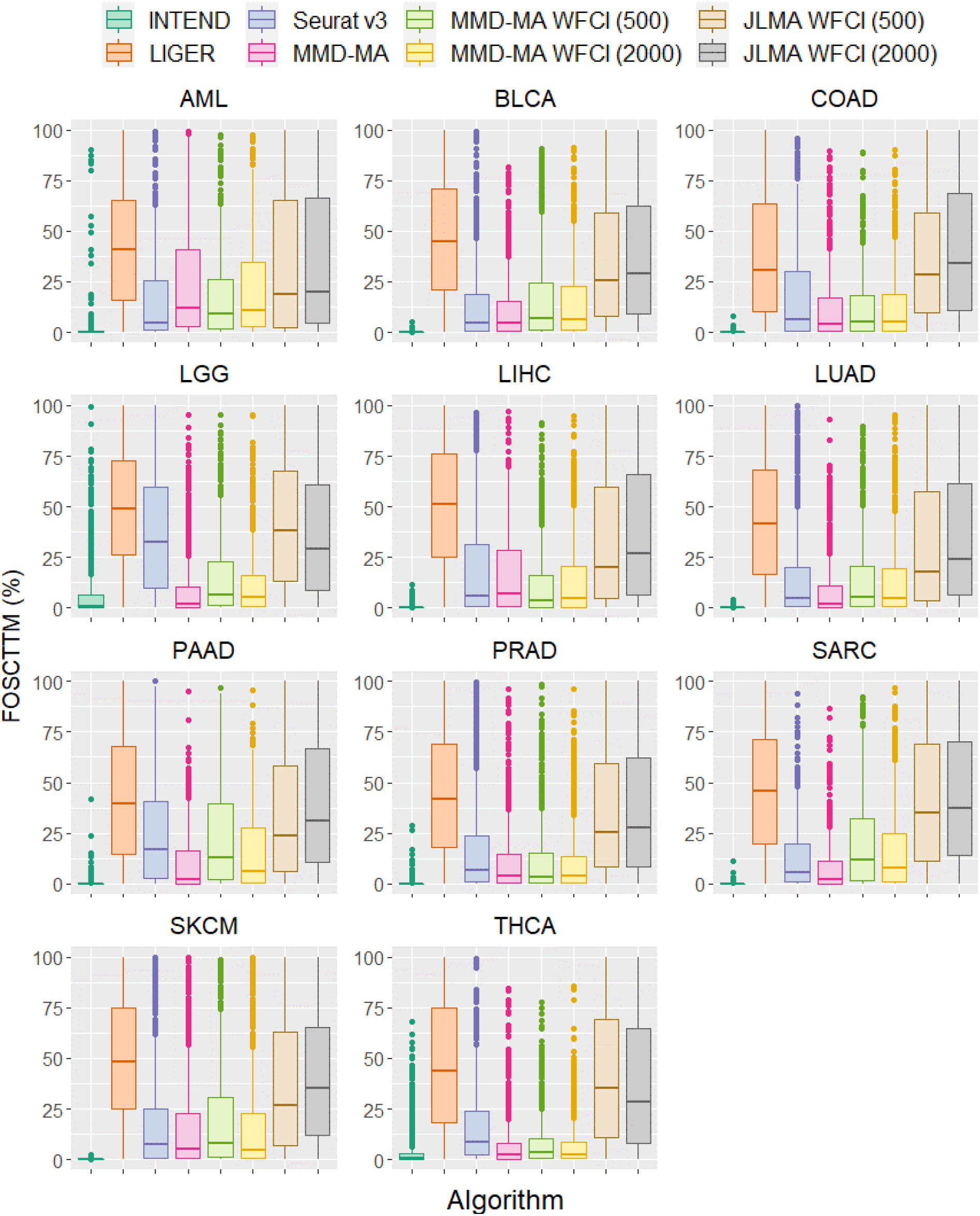
Distribution of FOSCTTM (%) scores in INTEND results on each cancer type.

We analyzed in more detail the results for the COAD dataset. We used UMAP (McInnes, Healy, and Melville 2018) for the 2D projection of the samples from the original omic feature spaces and from the integration shared space. **Figure 3** shows the results for INTEND, LIGER, Seurat, and MMD-MA algorithms. The results for JLMA WFCI and MMD-MA WFCI versions are presented in **Supplementary Figure 6**.

**Figure 3.**
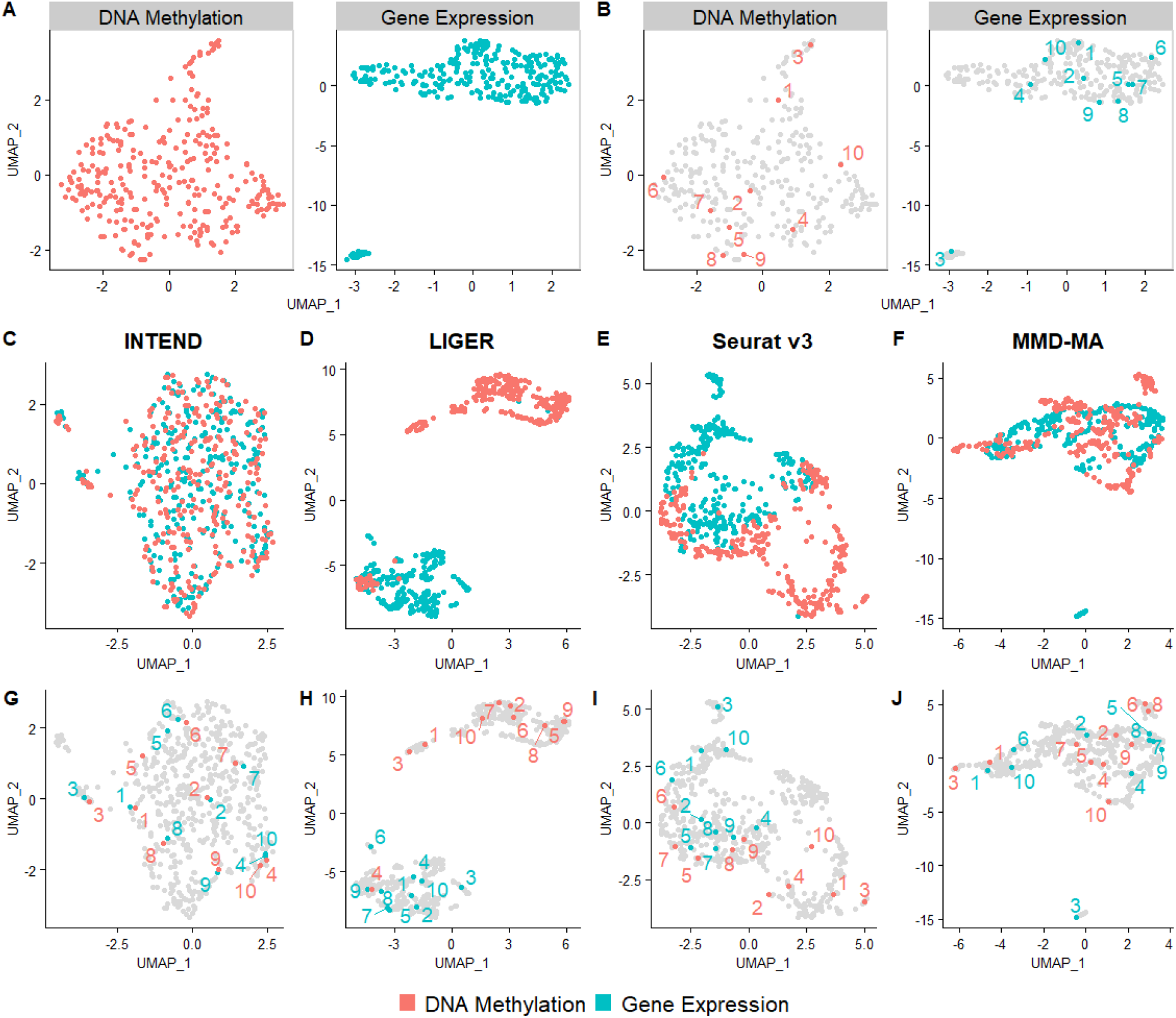
Results of integration of GE and DM samples from the colon adenocarcinoma dataset by different algorithms. (A) UMAP plots of the original data. (B) The same plots as in A. To appreciate concordance between omics, ten samples were randomly selected, and their matching points in both omics were labeled. (C-F) UMAP plots of the samples after they were projected to a shared space by each algorithm. (G-J) The same plots as in C-F with the selected points labeled. In all plots colors correspond to omics.

**Figure 3A-B** show the projections from the original feature spaces. One can appreciate that pairwise distances are not preserved between the omics. **Figure 3C-F** show for each algorithm the projections from the shared feature space. It is evident that the level of mixing between the two omics is highest for INTEND, intermediate for MMD-MA and lower for Seurat and LIGER. **3G-J** show the same projections as in **Figures 3C-F** with the 10 samples of **Figure 3B** marked. Evidently, INTEND does a much better job in projecting omics from the same sample to close positions. For example, the two points labeled 3 belong to distinct clusters of samples in both the DM and the GE spaces. INTEND was the only method to succeed in projecting the points from both omics into the same cluster in the shared space. A similar advantage of INTEND was obtained for all other cancer types, even when the average FOSCTTM was higher (**Supplementary Figures 7-16**).

### 3.2 Joint integration of multiple cancer types

In a second test, we applied the algorithms on four cancer datasets simultaneously. We used the datasets of COAD, LIHC, SARC and SKCM, covering 1448 GE and DM profiles. We did not supply the cancer type of each sample to the algorithms. We used the remaining seven TCGA datasets as the training set in INTEND’s training phase. INTEND performed this task with the best FOSCTTM integration score (**Supplementary Figure 17**), with perfect FOSCTTM for > 65% of the samples, and 1-2 orders of magnitudes better than the other methods: The mean scores were 0.37% for INTEND, 41.59% for LIGER, 9.33% for Seurat, and 4.01% for MMD-MA.

**Figure 4** shows 2D projections of the mapping by each of the algorithms. INTEND, Seurat and MMD-MA projected the samples from the different cancer datasets into separate clusters in the shared space (**Figure 4G-J**). In contrast, LIGER failed to preserve the biological variance among the tissue types, mapping samples of different types to the same clusters (**Figure 4I**). While INTEND mixed the samples from both omics in each cancer type cluster, Seurat and MMD-MA created clusters with substantial separation between the samples from each omic (**Figure 4C-F**).

**Figure 4.**
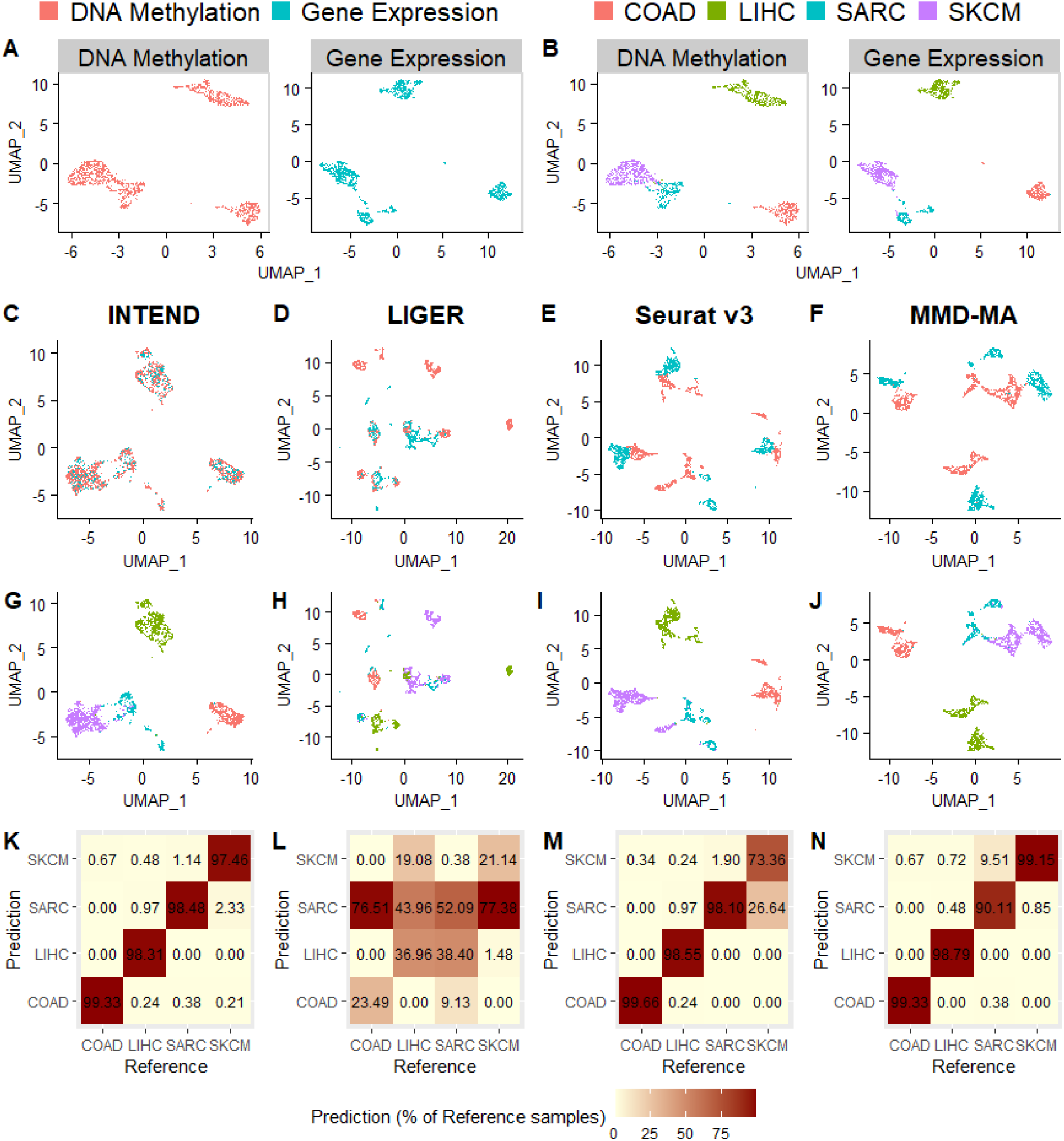
Results of joint integration of GE and DM samples of four cancer datasets: COAD, LIHC, SARC, and SKCM. (A-B) UMAP plots of the original data colored by omic (A) and by cancer type (B). (C-J) UMAP plots of the sample projections into the shared space by INTEND, LIGER, Seurat v3, and MMD-MA, colored by omic (C-F) and by cancer type (G-J). (K-N) Confusion matrices for the classification of the DM sample projections into cancer types based on majority vote among the five nearest GE samples in the shared space.

To further evaluate the results, we tested the quality of classifying the DM samples to specific cancer types based on the types of their neighboring GE samples in the shared space, as follows. Each DM sample was assigned by majority voting to the cancer type most represented among its five closest GE samples in the shared space. The confusion matrices between the inferred and true assignments are shown in **Figure 4K-N**. INTEND performed best, with > 97% of DM samples in each cancer type correctly classified. MMD-MA performed slightly worse: three cancer types had high accuracy classification, but the SARC cancer type had > 9% of the samples misclassified as SKCM. For Seurat, three cancer types had high accuracy classification, but the SKCM cancer type had >26% of the samples misclassified as SARC. The LIGER projections led to the lowest accuracy classification.

### 3.3 Using INTEND to identify subtypes in skin cutaneous melanoma

Clustering of single-omic cancer data is commonly used to identify subtypes. The quality of the clustering solution can be evaluated by the significance of separation in survival among subtypes. It has been observed that for certain cancer types, one omic may produce much better clustering than another. For example, Rappoport and Shamir (2018) benchmarked eight clustering algorithms on the TCGA SKCM data, and observed that GE profile clustering produced clusters with significant difference in survival in all algorithms, while in DM profile clustering only one algorithm showed such result. We hypothesized that in such cases, we could use INTEND to obtain GE predictions from the DM data, then jointly embed in the shared space the predictions and a set of GE profiles from the same cancer subtype, and achieve higher significance of separation in survival between clusters of the embedded predictions.

We used a dataset of 473 SKCM samples from TCGA that had both GE and DM profiles. We created 30 random partitions of this set into two equal disjoint groups, and for each partition, we used the first group’s DM profiles and the second’s GE profiles. We used INTEND to obtain a predicted gene expression matrix (P) from the DM samples and then embed P jointly with the GE profiles. Call the embedded P data EP. For the training phase of INTEND model, we used samples from all TCGA datasets listed in **Table 1** but excluded the SKCM dataset.

We first clustered separately the original partitioned DM and GE data. We performed each clustering task using k-means (see Methods) after selecting the 2000 features with the highest variance and normalizing the features to have zero mean and a standard deviation of one (as in Rappoport and Shamir (2018)). We ran the algorithm for *k* between 2 and 15, and selected the desired number of clusters using the “elbow method” (see Methods). We measured differential survival between clusters by computing the p-value for the log-rank test. We estimated the p-values using permutation tests (Rappoport and Shamir 2019a). As we hypothesized, in most cases, the clustering of the GE data obtained more significant differential survival between clusters than the clustering of the DM data, with the log-rank p-value of the first being lower in 27 of the 30 partitions.

Next, for each of the 30 partitions, we used INTEND’s joint embedding of the DM and GE samples to classify the DM samples based on the k-means clustering of the GE samples. Each DM sample was assigned by majority voting (with ties broken at random) to the cluster most represented among the five GE embeddings closest to its matching EP representation in the shared space. In 23 of the 30 splits, clustering the DM samples using this method obtained more significant differential survival than using the k-means clustering of the DM samples. The average log-rank p-values for the clusterings for all 30 random splits were: 0.07 for the GE k-means clustering, 0.56 for the DM k-means clustering, and 0.21 for the integration-based DM clustering, as described above.

We further investigated one of the 23 partitions for which the integration-based DM clustering achieved more significant differential survival than the DM clustering. For that partition, the DM clustering resulted in two clusters with insignificant differential survival (p-value=0.978, **Figure 5A**), whereas the GE clustering resulted in two clusters with significant differential survival (p-value=0.018, **Figure 5B**). The integration-based DM clustering also obtained significant differential survival between clusters (p-value=0.036, **Figure 5C**).

**Figure 5.**
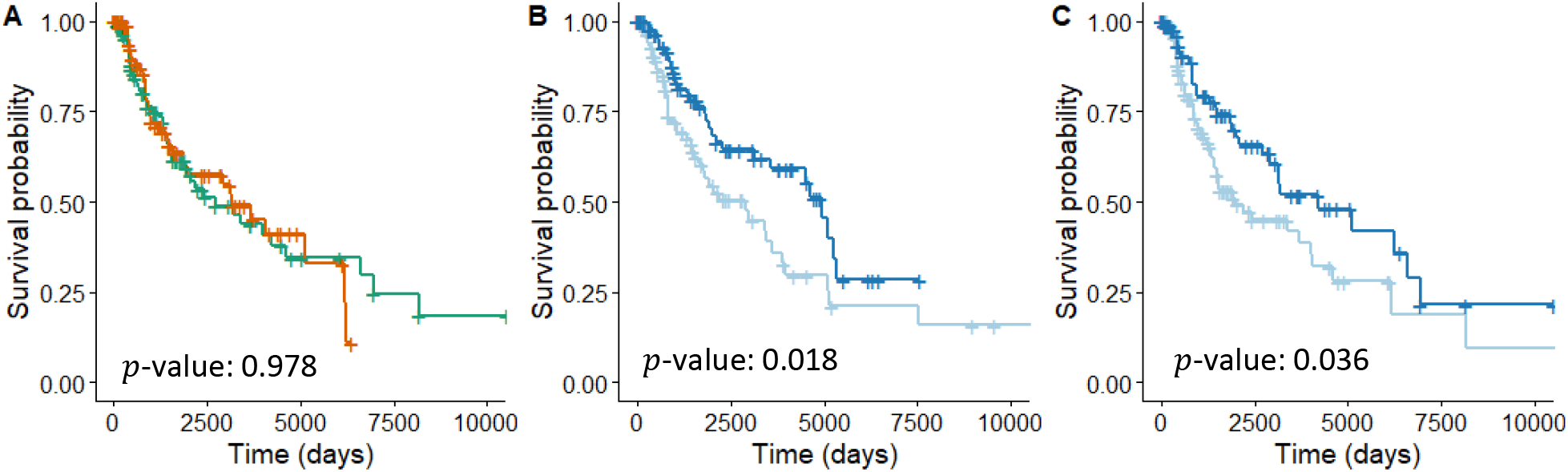
Kaplan-Meier plots of clusters of SKCM patients obtained using DM profiles, GE profiles, and their INTEND embeddings. (A) Plot for clusters of the original DM profiles. (B) Plot for clusters of the original GE profiles. (C) Plot for clusters of the DM profiles obtained by the integration-based clustering. See **Supplementary Figure 18A-E** for the UMAP plots and the clusters.

Next, we tested whether the subtypes obtained by the integration-based DM clustering were biologically or clinically more similar to those obtained by the GE k-means clustering. We found that primary tumor and metastases samples were represented in each of the DM k-means clusters exactly in their portion of all DM samples (18.26% of primary tumor samples in both clusters). By contrast, when looking at the GE clusters, the primary tumor samples were overrepresented in one cluster and underrepresented in the other (28.21% of primary tumor samples in the first cluster, 5.94% in the second, 17.89% in all GE samples). We observed a similar pattern in the integration-based DM clustering: 23.77% of primary tumor samples in one cluster and 11.34% in the other (and 18.26% in all DM samples). This example shows the potential of transferring biological information between GE and DM samples measured on different populations, using INTEND’s integration.

### 3.4 Joint analysis of lung adenocarcinoma datasets from different sources

Our next goal was to test the utility of INTEND in joint analysis of two datasets, one of DM profiles and one of GE profiles, coming from different sources. We used data from two studies of LUAD: GE of 172 tumor samples from Chen et al. (2020), and DM profiles of 477 samples from TCGA. The datasets were collected in different studies covering disjoint groups of LUAD patients.

#### 3.4.1 Integration

For the training phase of the model, we used samples from all TCGA datasets listed in **Table 1** but excluded the LUAD dataset. The integration results are summarized in **Figure 6A-B.** As the two target datasets here are disjoint we cannot use FOSCTTM to evaluate their mixing in the embedding phase. As a sanity check, we considered for each sample its closest 32 neighbors (5% of the samples) in the shared space. We expected that if the local neighborhood of a sample is well mixed, the number of samples from each omic in the neighborhood would reflect the relative sizes of the target datasets. For each sample we measured the ratio between the observed and expected number of samples from the other omic in its neighborhood. If the omics are fully separated we would expect this ratio to be near zero, whereas for perfectly mixed samples we would expect it to be close to 1. The mean computed ratio for all samples in the shared space was 1.003 (*SD* = ±0.258), and the *IQR* was 0.82 – 1.15, indicating well-mixed samples across omics.

**Figure 6.**
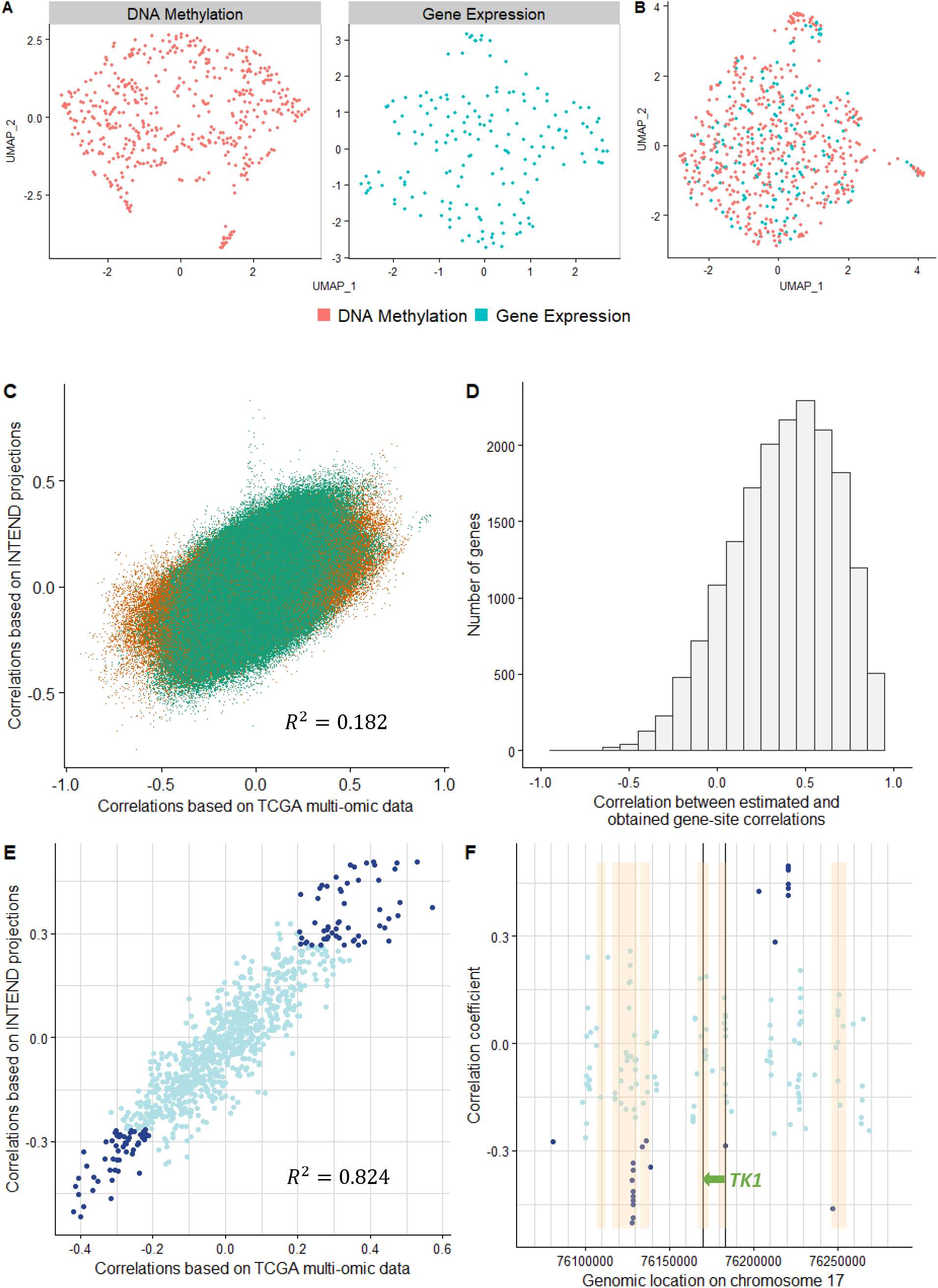
INTEND results on LUAD GE profiles from Chen et al. (2020) and DM profiles from TCGA. (A-B) UMAP plots of the original data (A) and of the projections into the shared space (B), colored by omic. (C) Scatterplot of the estimated correlations based on the matching of INTEND projections versus the observed correlations from the multi-omic TCGA dataset, for each of the considered 10.14 million gene-site pairs. The pairs for which the site is within 10Kb from the gene are colored in orange. These gene-site pairs were considered in INTEND training phase on the TCGA datasets (excluding LUAD). (D) Histogram of the correlation between the estimated and TCGA-observed gene-site correlations, per gene. (E) Correlation coefficients between TK1 expression and methylation levels, at 964 sites located ±1Mb from TK1. Y axis: correlations when TK1 expression is based on INTEND projections; x axis: correlations when both the GE and the paired DM data were taken from TCGA. Correlations with ***p***-value < **1**e-5 based on both methods are colored in dark blue (F) Estimated correlation coefficients based on INTEND projections in sites located ±100Kb from TK1. The x axis shows their genomic location (build GRCh37/hg19). Correlations with ***p***-value < **1**e-5 are colored in dark blue, TK1 location is marked by the green arrow. The highlighted yellow regions indicate enhancer regions supported by at least four GeneHancer sources

#### 3.4.2 Correlations between methylation at specific sites and expression

Next, we wished to test if INTEND application on the two datasets can be used to reveal connections between specific distal DM sites and the regulation of GE in LUAD tumors, even though the GE profiles and DM profiles used here were collected from disjoint sets of patients. For this task, we extracted the estimated correlations between methylation levels at specific CpG sites and the expression levels of specified genes as follows.

We considered for every gene *g*, the methylation sites located within ±1Mb of *g* (including sites in *g*). There was a total of approximately 10.14 million such gene-site pairs, for which the expression and methylation levels were measured, covering 18,553 different genes. Recall that INTEND model was trained using proximal sites located only within ±10Kb from each gene, while here we explore mostly distal methylation sites. To estimate the correlation between the methylation level at site s and the expression level of gene *g*, we used INTEND projections to get matchings between GE and DM profiles from different patients. First, to match GE and DM profiles, we found the mutual nearest neighbors between the projections of all DM and GE samples in the shared space, using the *batchelor* R package (Haghverdi et al. 2018). A pair of a GE profile *e* and a DM profile m was considered a match if the projection of m was in the *k*-nearest neighbors of the projection of *e* and vice versa (i.e. the projections of *e* and *m* are mutual *k*-nearest neighbors). For *k* = 5 we obtained 270 matches between GE and DM profiles (out of 172 · 477 = 82,044 possible matches). Next, using the 270 matches, we computed the Pearson’s correlation coefficient and tested the statistical significance of the association between the expression level and the methylation level of each considered gene-site pair.

We wished to assess the validity of the estimated correlations, based on the created 270 matchings of GE and DM samples from the two LUAD datasets (from here on: “estimated correlations”). We compared the estimations to the correlations obtained from 477 pairs of GE and DM profiles measured from the same tissue, from the multi-omic LUAD TCGA dataset. For each of the approximately 10.14 million gene-site pairs previously described, we also computed the correlation between the expression of the gene and the methylation level of the relevant site, based on the multi-omic TCGA dataset (from here on: “TCGA-observed correlations”). **Figure 6C** shows for each gene-site pair the estimated correlation versus the TCGA-observed correlation. Approximately 5.08% of the considered gene-site pairs were detected with significant correlation (*p*-value< 0.01), either positive or negative, according to both methods. For 95.63% of these significant pairs, the estimated correlation coefficient had the same sign as the TCGA-observed correlation. We also tested for each of the considered genes, the correlation between the estimated correlation and the TCGA-observed correlation, for all sites relevant for that gene. Out of the 18,553 considered genes, there was a significant positive correlation between the estimated and TCGA-observed correlations (p-value < 0.05) for 14,693 of the genes. The correlation between the estimated and TCGA-observed correlations was above 0.8 for 1,041 of the genes, and above 0.9 for 180 of them **(Figure 6D**). This demonstrates the potential of INTEND integration method to uncover connections between DNA methylation and the regulation of gene expression, both for proximal and distal methylation sites.

##### 3.4.2.1 An in-depth look at the regulation of Thymidine Kinase 1

We chose to look in detail at the distal methylation sites of the gene Thymidine Kinase 1 (TK1). High expression of TK1 was recorded in many solid tumors, and was associated specifically with poor prognosis of patients with LUAD (Malvi et al. 2019; Jagarlamudi and Shaw 2018; He et al. 2010). We computed the correlation between the methylation levels in 964 sites within ±1Mb from TK1, and its expression level. The estimated correlations based on the matching of GE and DM profiles from INTEND projections were highly concordant with the correlations computed using the multi-omic TCGA dataset (*R*^2^ = 0.824, **Figure 6E**).

Methylation patterns in enhancer regions are known to be altered in cancer and are closely linked to changes in expression of cancer-related genes (Aran, Sabato, and Hellman 2013). Therefore, we checked if strong expression-methylation correlations extracted from INTEND projections can indicate potential distal enhancer regions. We used the GeneHancer database of enhancers and their inferred target genes (Fishilevich et al. 2017) for information on TK1 enhancers. There were eight enhancer regions supported by at least four GH sources, seven of them within a 100Kb range from TK1. **Figure 6F** shows the enhancer regions located ±100Kb from TK1, and the correlations between methylation and TK1 expression, for sites located in this range. 14 out of the 15 sites in this range with strong negative correlation (p-value < 1e-5), are located in one of the documented enhancer regions. Note that all but two of them fall outside the ±10Kb used for the training phase.

Out of the 964 sites in 1Mb range from TK1, we investigated the ten sites with the strongest negative estimated correlations (full details in **Supplementary Table 2**). Eight of them are located in two of the enhancer regions shown in **Figure 6F** (seven of them in a short interval of less than 500 bases). The other two sites, cg11868461 and cg05110391, are located approximately 350Kb downstream and 400kB upstream the TSS, respectively. They were not in one of the regions marked by GeneHancer as TK1 enhancers. Nevertheless, both cg11868461 and cg05110391 were identified as “enhancer probes” (not specifically related to TK1) by Mullen et al. (2020), using H3K27ac ChIP-seq data from normal and tumor lung tissue samples to identify lung-relevant enhancer regions.

## 4 Discussion

We presented the INTEND algorithm for integrating gene expression and DNA methylation from different datasets. We tested it on multiple multi-omic cancer datasets and compared it with extant multi-omic integration algorithms. INTEND showed significantly superior results on all tested datasets when integrating data from single and multiple cancer types, both in terms of FOSCTTM score and in classification to cancer types according to the integration results. We demonstrated the potential of INTEND to transfer biological information between GE and DM samples measured on non-overlapping populations of skin cutaneous melanoma patients. Clustering DM samples achieved higher significance of separation in survival between clusters when using the integration results of the DM and GE data, than using the original DM data only. In another typical use case, we tested INTEND in joint analysis of two lung adenocarcinoma datasets from different sources. Here INTEND demonstrated its potential to uncover connections between DNA methylation and the regulation of gene expression.

INTEND’s novelty mainly resides in the incorporation of the prediction of a GE profile from a DM profile of a sample, into the MO/MD integration problem. Unlike algorithms such as LIGER and Seurat, which were developed mainly to solve the SO/MD problem and then were extended to solve the MO/MD problem, INTEND suggests another method to generate the correspondence information between features – a paramount part for the integration. INTEND presents a data-driven approach to generate a predicted gene expression matrix, thus effectively reducing the MO/MD problem of integrating GE and DM profiles to the simpler SO/MD problem of integrating multiple GE datasets. Importantly, the data necessary for the training phase of INTEND can represent different populations than the data used for the embedding phase. In all cases presented in this paper, the used training data originated from samples from other cancer types than represented in the target datasets for integration.

INTEND has several limitations. First, the training phase requires multi-omic data measured on the same set of samples, which is not required for the other algorithms we tested. While the training data is not required to be from a similar population to the target data, it is necessary that the omics will be measured in the same method on the train and target datasets. Obtaining multi-omic measurements may be harder in several scenarios, e.g. single-cell multi-omic data. Second, the final step in the embedding, applying CCA, may be less effective when the target datasets contain non-overlapping sample populations (e.g. when one of the target datasets contains a group of samples from a cancer type which is not present in the second). Stuart et al. 2019 addressed this limitation of using CCA as a final step and introduced a method to overcome it, using the concept of mutual nearest neighbors to identify anchors between the target datasets.

Lastly, we note two possible directions of extending this work. The first is the integration of other pairs of omics, in addition to GE and DM, in a similar method. Here we used an established, simple biological observation, namely the relation between the state of proximal methylation sites to the gene’s expression, to build a model and uncover the connections between GE and DM based on available multi-omic data. This concept may be extended to other pairs of omics with available data measuring both on the same set of samples. Another future research direction is the incorporation of methods from algorithms tackling the SO/MD integration problem, after the first step in INTEND’s embedding phase, which results in the predicted GE matrix.

## Acknowledgments

The results published here are in whole or part based upon data generated by the TCGA Research Network: https://www.cancer.gov/tcga. Study supported in part by Israel Science Foundation (grant No. 3165/19, within the Israel Precision Medicine Partnership program, and grant No. 1339/18), German-Israeli Project DFG RE 4193/1-1, and the Raymond and Beverly Sackler Chair in Bioinformatics, Tel Aviv University. NR was supported in part by a fellowship from the Edmond J. Safra Center for Bioinformatics at Tel-Aviv University, by the Planning and Budgeting Committee (PBC) fellowship for excellent PhD students in Data Sciences, by the Tel Aviv University Healthy Longevity Research Center, and by the Herczeg Institute on Aging.

