## Supplementary Information for "Integration of Gene Expression and DNA Methylation Data Across Different Experiments"

### Data Collection

The genes' coordinates used in the training phase were taken from the GRCh37/hg19 assembly (Church et al. 2011). The data was downloaded from the UCSC Genome Browser (Haeussler et al. 2019) from the following URL: https://hgdownload.soe.ucsc.edu/goldenPath/hg19/bigZips/genes/hg19.refGene.gtf.gz. Build version 37 of the assembly was used to match the genome coordinates in the methylation probes manifest file.

The methylation data we used was measured using Infinium HumanMethylation450 v1.2 BeadChip by Illumina. The genome mapping information of the methylation probes was downloaded from Illumina's website (ftp://webdata2:webdata2@ussd-ftp.illumina.com/downloads/ProductFiles/HumanMethylation450/HumanMethylation450_15017482_v1-2.csv).

The TCGA data was downloaded using the TCGA-Assembler software (Wei et al. 2018; Zhu, Qiu, and Ji 2014). The DNA methylation data was downloaded using the 'DownloadMethylationData' function. The RNA-seq data was downloaded with the 'DownloadRNASeqData' function, setting the 'assayPlatform' parameter to 'gene.normalized_RNAseq'.

The additional LUAD dataset (Chen et al. 2020) was downloaded from OncoSG, the Singapore Oncology Data Portal (https://src.gisapps.org/OncoSG/), under 'Lung Adenocarcinoma (GIS, 2019)'.

### Correspondence information between features

Some of the tested multi-omic integration algorithms require correspondence information between the features across omics. LIGER and Seurat assume that the input matrices to be integrated share the same set of features. When these methods were previously used to integrate scRNA-seq and methylation (LIGER) or scATAC-seq (Seurat) data, the input from the latter omic was converted to a matrix with gene-level features. The new features were expected to correspond to the GE features.

To summarize gene-level methylation, we used the annotations of methylation sites into six possible regions: TSS1500 (201-1500 bps upstream of the transcription start site(TSS)), TSS200 (0-200 bps upstream of the TSS), 5'UTR (untranslated region), 1stExon, Body, and 3'UTR. The HumanMethylation450 BeadChip annotations were taken from Ilumina (https://support.illumina.com/downloads/humanmethylation450_15017482_v1-2_product_files.html). Of those, we used the sites in the TSS1500, TSS200, 5'UTR and 1stExon regions. We chose these regions as they showed anticorrelation with GE on the TCGA data (**Supplementary Figure 4**). This matched previous reports on anti-correlation between DM levels in the promoter region and the gene's expression level (Deaton and Bird 2011). The final gene-level summary was minus the average methylation signal in those regions.

### Overview of the Methods Used

#### *LIGER*

LIGER (Welch et al. 2019) takes as input multiple single-cell datasets, either scRNA-seq experiments or multi-omic measurements. In the latter case, LIGER takes as input the preprocessed datasets after conversion to a shared gene-level feature space. When LIGER is used to integrate gene expression with methylation data from mouse frontal cortical neurons, the methylation data is first converted to gene-level methylation features (non-CpG gene body methylation). The direction of the methylation signal is reversed to incorporate the assumption of general anti-correlation with gene expression in neurons (Mo et al. 2015). LIGER then employs integrative non-negative matrix factorization to create for each matrix a dataset-specific factor plus a shared factor across the datasets. The shared factor is used to jointly embed cells in a common low-dimensional space.

*Seurat v3*

The Seurat v3 algorithm (Stuart et al. 2019) was designed to integrate multiple scRNA-seq datasets in the SO/MD setting, but was also demonstrated to integrate scATAC-seq and scRNA-seq data in the MO/MD setting. The first step of such integration is similar to LIGER. The scATAC-seq data is converted to a gene-activity matrix, based on the accessibility of sites proximal to the gene's transcription start site (Pliner et al. 2018). The gene activity matrix has the same feature set as the scRNA-seq matrices and it is assumed to be correlated with them. Seurat first uses canonical-correlation analysis to jointly reduce the dimension of the two datasets to a shared space. Then it identifies mutual nearest neighbors across the datasets. The pairings found are termed "anchors". These anchor pairs are scored based on the consistency of anchors across the neighborhood structure of each dataset. The scored anchors are utilized to compute a projection mapping for each cell to embed it in the shared space.

*JLMA*

The joint Laplacian manifold alignment (JLMA) algorithm (C. Wang and Mahadevan 2008) learns a projection that maps datasets from two different feature spaces to a shared lower-dimensional space. This is done while simultaneously preserving the neighborhood relationships in each set and matching the local geometry of samples from the two sets. JLMA constructs a joint Laplacian matrix across the two domains, which captures the similarities within each dataset and the similarities across the datasets. The similarities across the datasets can be given as input to the algorithm (in a semi-supervised manner) or computed by the algorithm according to matching between the local geometry of the samples. In the latter case, JLMA does not require any correspondence information. The local geometry measure is computed based on the $k$-NN graph of each dataset. Finding the local geometry matching is computationally expensive even for small values of $k$, as it runs in $O(k!)$ time. After the joint Laplacian is computed, the optimal solution is found by solving a generalized eigenvalue decomposition problem.

*MMD-MA*

The maximum mean discrepancy-manifold alignment (MMD-MA) algorithm (Liu et al. 2019) is an unsupervised manifold alignment algorithm. It was created specifically for the task of single-cell multi-omics integration. The algorithm assumes the samples from the different omic datasets are drawn from the same initial population, but it does not require any correspondence information between the samples or the features. MMD-MA seeks an optimal alignment by minimizing an objective function with three terms. The first is the maximum mean discrepancy, which corresponds to the distance between the two mapped manifolds in the shared space. The second term, named distortion, measures relationships among data points between the original space and the shared latent space. The third is a penalty term that is intended to avoid a collapse to a trivial solution.

### Benchmark Methods and Software

All experiments ran on R-4.0.1 (and python 3.8.0 for MMD-MA). For all methods, we followed the usage guidelines supplied by the creators. The preprocessing steps described in section 2.2.3 were applied to the input GE and DM data for all algorithms used in the benchmark.

*LIGER*
**Implementation:** We used the 'rliger' R package, version 0.5.0. The methods referred to in the following subsection were supplied by this package. We followed LIGER guidelines for integrating GE and DM data (https://welch-lab.github.io/liger/rna-methylation.html).

**Preprocessing**: No further preprocessing (except the steps described in 2.2.3) was applied to the input GE and DM matrices. The aggregated gene-level methylation (described above in this supplement) was used as the methylation input for LIGER. The GE data was normalized and scaled using the `normalize` and `scaleNotCenter` methods. The DM input was not normalized and scaled as suggested by the guidelines.

**Feature selection**: The genes were selected using the `selectGenes` method when considering only the GE data, as suggested by the guidelines. This method selects the variable genes by comparing the variance of each gene's expression to its mean expression.

**Execution details**: The default settings were used in the factorization and quantile normalizations phases of LIGER. As suggested by the guidelines, the `quantileAlignSNF` method was used with `center=T`, considering the density of the methylation data. The factorization was done with k (the number of factors) between 2 and 40, resulting in data projections in 2 to 40 dimensions.

*Seurat v3*
**Implementation:** We used the 'Seurat' R package, version 3.2.3. The methods referred to in the following subsection were supplied by this package. We followed Seurat guidelines for integration and label transfer (https://satijalab.org/seurat/archive/v3.2/integration.html).

**Preprocessing**: No further preprocessing (except the steps described in 2.2.3) was applied to the input GE and DM matrices. The aggregated gene-level methylation (described above in this supplement) was used as the methylation input for Seurat. The GE data was normalized using the `NormalizeData` method with the relative count normalization method. This step is not documented in the guidelines but empirically improved the results in all tested cases.

**Feature selection**: The genes were selected using the `FindVariableFeatures` method with the default parameters and selection method.

**Execution details**: The default settings were used. In the step of identifying anchors (using `FindIntegrationAnchors`), we used 30 neighbors when filtering the anchors (`k.filter=30`). We ran the algorithm with all possible dimensions between 2 and 40.

*JLMA*
**Implementation:** We used our implementation based on the JLMA paper, as we didn't find an R package implementing JLMA. The implementation code is part of the INTEND project on GitHub.

**Preprocessing**: No further preprocessing (except the steps described in 2.2.3) was applied to the input GE and DM matrices. The aggregated gene-level methylation (described above in this supplement) was used as the methylation input for JLMA.

**Feature selection**: We selected the $n$ genes with the highest variance in expression for $n = 500$ and 2000 (the algorithm ran in two variants). We scaled the inputs such that each feature (gene) had zero mean and unit variance.

**Execution details**: As mentioned in subsection 2.2.4, we computed the cross-omic similarity matrix for JLMA based on the aggregated gene-level methylation matrix. We used the hyper-parameter $\mu = 1$.

*MMD-MA*

**Implementation:** The algorithm's source code was downloaded from https://noble.gs.washington.edu/proj/mmd-ma/. We made minor changes in the source code to allow us to run the algorithm for the desired dimensions.

**Preprocessing**: No further preprocessing (except the steps described in 2.2.3) was applied to the input GE and DM matrices. As mentioned in subsection 2.2.4, we ran MMD-MA with both the original methylation data and the aggregated gene-level methylation (described above in this supplement) as inputs.

**Feature selection**: When using the original methylation data, no feature selection method was applied before computing the inter-similarity matrices for the GE and DM inputs. When running MMD-MA with the gene-level methylation data, we selected the $n$ genes with the highest variance in expression for $n =$ 500 and 2000 (the algorithm ran in two variants). In this case, we scaled the inputs such that each feature (gene) had zero mean and unit variance.

**Execution details**: We ran MMD-MA with dimensions 2,10,20,30, 40. We did not run it for all possible dimensions between 2 and 40 due to long running times.

### CCA optimization problem solution

To solve the optimization problem in equation (3), we use the Lagrange multipliers method. We denote $K = X^T Y$. Let:

$$L = u^T K v - \frac{\lambda_1}{2}(u^T u - 1) - \frac{\lambda_2}{2}(v^T v - 1) \tag{5}$$

Differentiating $L$ with respect to $u$ and $v$ gives:

$$\frac{\delta L}{\delta u} = Kv - \lambda_1 u = 0 \rightarrow \; Kv = \lambda_1 u \tag{6}$$

$$\frac{\delta L}{\delta v} = K^T u - \lambda_2 v = 0 \rightarrow K^T u = \lambda_2 v \tag{7}$$

Left-multiplying (6) and (7) by $u^T$ and $v^T$ respectively, and using the constraints $||u||_2^2 = 1$ and $||v||_2^2 = 1$:

$$\lambda_1 = u^T K v = v^T K^T u = \lambda_2 \tag{8}$$

Thus $u$ and $v$ are the left and right unit singular vectors of $K$ with singular value $\lambda = \lambda_1 = \lambda_2$. Since the objective is to maximize $u^T K v$, then $u_1, v_1$ are the left and right unit singular vectors of $K$ with the greatest singular value. We claim that $\forall i \in \{1, \dots, d\}, u_i$ and $v_i$ are the left and right unit singular vectors of $K$ with the $i^{th}$ greatest singular value. Let $u_i$ and $v_i$ be the $i^{th}$ unit singular vectors of $K$. Then we showed that $(u_i, v_i)$ maximizes over all $u \in \mathbb{R}^{n_E}, v \in \mathbb{R}^{n_M}$, the correlation between $Xu$ and $Yv$. As $u_i^T u_j = v_i^T v_j = 0$ for $j < i$, and we assumed that $X^T X$ and $Y^T Y$ are diagonal, then $Cor(Xu_i, Xu_j) = Cor(Yv_i, Yv_j) = 0$ for $j < i$. Hence the optimal canonical-correlation vectors can be obtained by SVD of $K = X^T Y$. We denote $U = (u_1, u_2, \dots, u_d) \in \mathbb{R}^{n_E \times d}$ and $V = (v_1, v_2, \dots, v_d) \in \mathbb{R}^{n_M \times d}$ where $u_i$ *and* $v_i$ are the $i$-th left and right singular vectors, respectively. The output of this step is the matrix $S = [U^T \quad V^T]$, of dimensions $d \times (n_E + n_M)$, containing the embeddings of samples from both target sets in the shared $d$-dimensional space.

### Supplementary Tables

**Supplementary Table 1.** Number of features and samples in each TCGA dataset before and after the handling of missing values

| Dataset | Number of samples | | | | Number of features | | | |
|---|---|---|---|---|---|---|---|---|
| | Gene expression | | DNA methylation | | Gene expression | | DNA methylation | |
| | Before | After | Before | After | Before | After | Before | After |
| **AML** | 173 | 173 | 194 | 194 | 20530 | 20530 | 526729 | 432429 |
| **BLCA** | 427 | 427 | 440 | 440 | 20530 | 20530 | 526729 | 431716 |
| **COAD** | 328 | 328 | 353 | 353 | 20530 | 20530 | 526729 | 431308 |
| **LGG** | 534 | 534 | 534 | 534 | 20530 | 20530 | 526729 | 431991 |
| **LIHC** | 424 | 424 | 430 | 430 | 20530 | 20530 | 526729 | 430791 |
| **LUAD** | 576 | 576 | 507 | 507 | 20530 | 20530 | 526729 | 431486 |
| **PAAD** | 183 | 183 | 195 | 195 | 20530 | 20530 | 526729 | 428806 |
| **PRAD** | 550 | 550 | 553 | 553 | 20530 | 20530 | 526729 | 432201 |
| **SARC** | 265 | 265 | 269 | 269 | 20530 | 20530 | 526729 | 428486 |
| **SKCM** | 473 | 473 | 475 | 475 | 20530 | 20530 | 526729 | 430579 |
| **THCA** | 572 | 572 | 571 | 571 | 20530 | 20530 | 526729 | 432307 |

**Supplementary Table 2.** Top ten methylation sites with the strongest negative estimated correlations out of the 964 sites in 1Mb range from TK1

| Methylation site | Location on chromosome 17 (build GRCh37/hg19) | Correlation coefficient estimation |
|---|---|---|
| cg11868461 | 75830800 | -0.5181234 |
| cg06643271 | 76128170 | -0.5016554 |
| cg24988684 | 76128556 | -0.4887382 |
| cg10460946 | 76247467 | -0.4631925 |
| cg11493223 | 76128522 | -0.4516062 |
| cg02911077 | 76128621 | -0.4396759 |
| cg18901278 | 76128531 | -0.4280529 |
| cg04947157 | 76128481 | -0.4135805 |
| cg03742808 | 76128634 | -0.4130168 |
| cg05110391 | 76588634 | -0.4063449 |

### Supplementary Figures

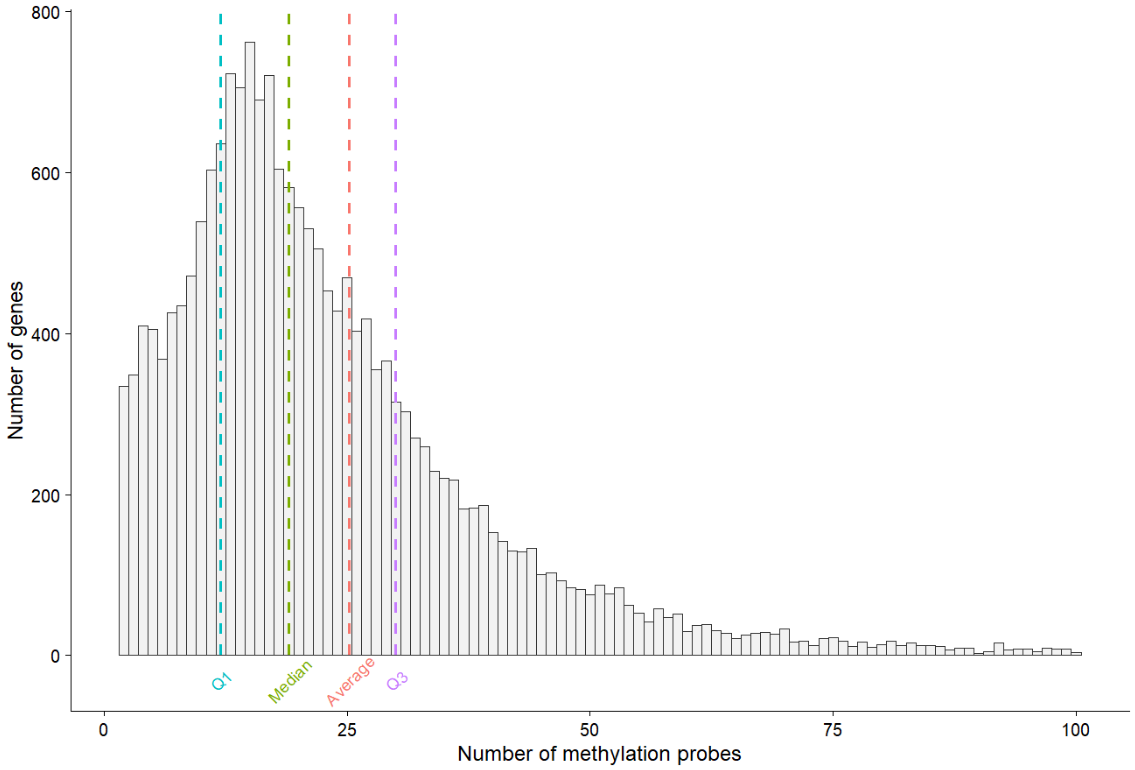


**Supplementary Figure 1**

Histogram of the number of methylation sites per gene. Average: 25.22, median: 19, interquantile range ($IQR$): 12-30. The maximum number of methylation sites per gene was 1055 (outside the plot axis limits).

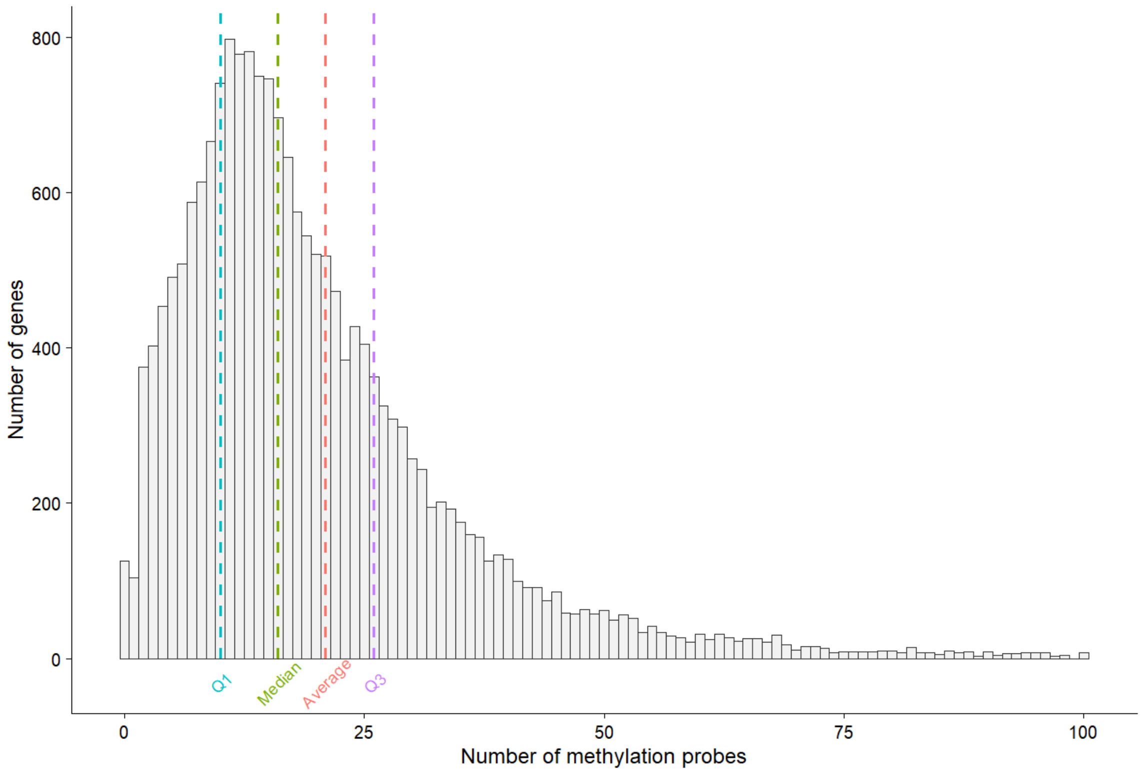


**Supplementary Figure 2**

Histogram of the number of methylation sites per gene in the model after Lasso shrinkage on the TCGA data. The model was trained on ten cancer subtypes data from TCGA: AML, BLCA, COAD, LGG, LIHC, PAAD, PRAD, SARC, SKCM, and THCA. Average: 20.93, median: 16, interquantile range ($IQR$): 10-26. The maximum number of methylation sites per gene was 424 (outside the plot axis limits).

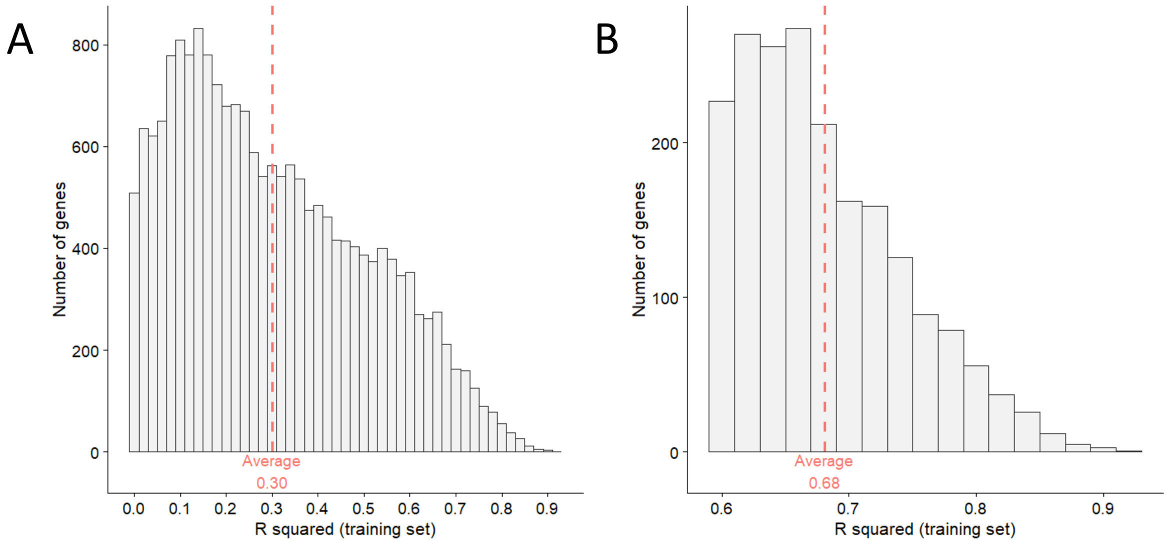


**Supplementary Figure 3**

Histograms of $R^2$ values between predicted and observed gene expression, when training on GE and DM data of 10 cancer subtypes from TCGA (the datasets listed in **Table 1**, excluding LUAD), covering 3852 tumor samples. (A) All 19143 genes, (B) The 2000 genes with the highest $R^2$.

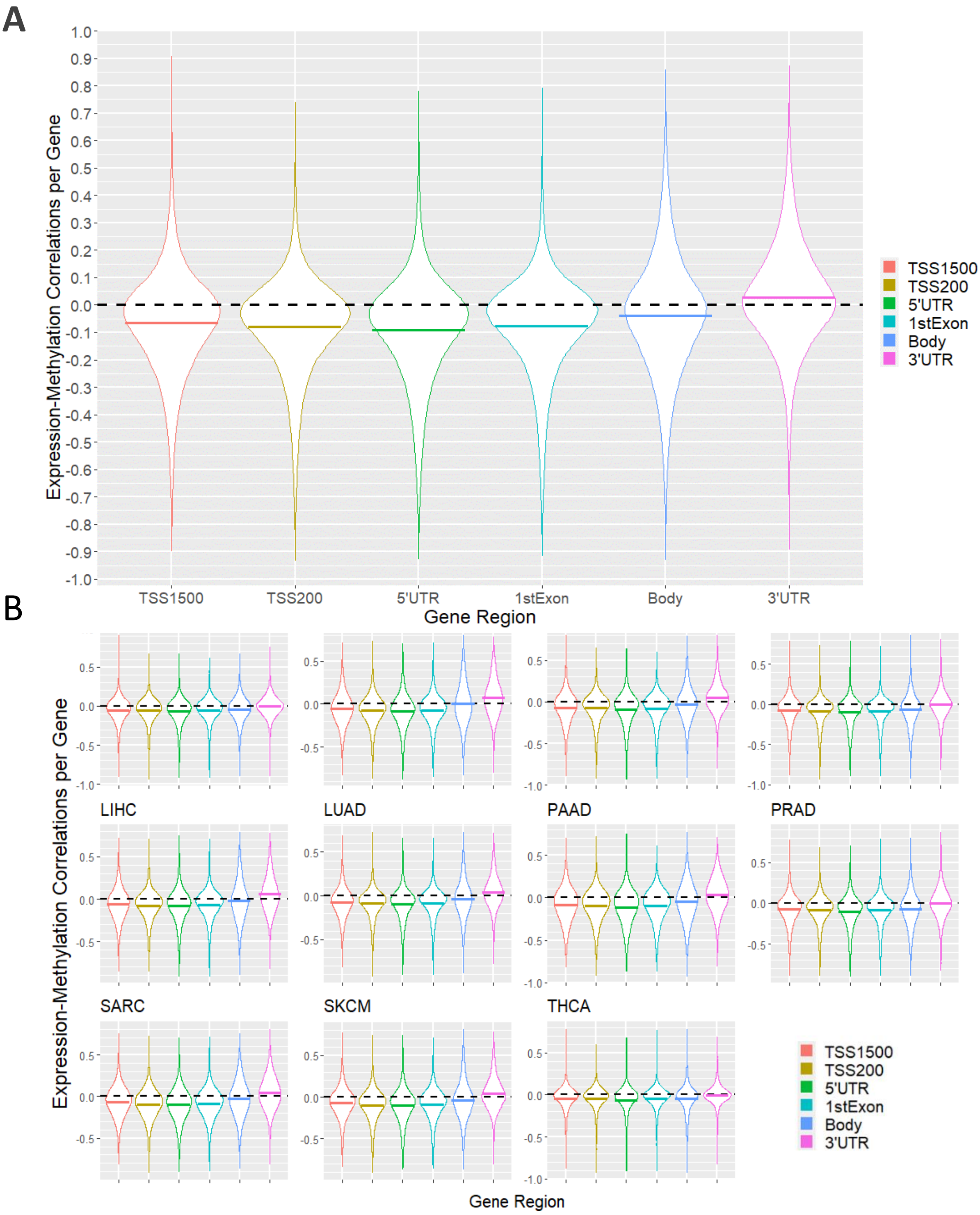


**Supplementary Figure 4: Correlation between gene expression and DNA methylation levels in different genomic regions**

Correlations were measured for genes that had both expression and methylation data in the specified region of the gene. Data included samples from eleven cancer types from the TCGA database (**Table 1**). The horizontal line in each violin plot is the mean correlation for the region, and the black dashed line shows a correlation of zero. (A) Summary over all subtypes, (B) Results for each subtype separately. The mean correlation for TSS1500, TSS200, 5'UTR, and 1stExon is $< -0.04$ for every subtype, and $< -0.065$ on average across subtypes. The Body regions exhibit a positive mean correlation for one subtype (BLCA), and 3'UTR for seven.

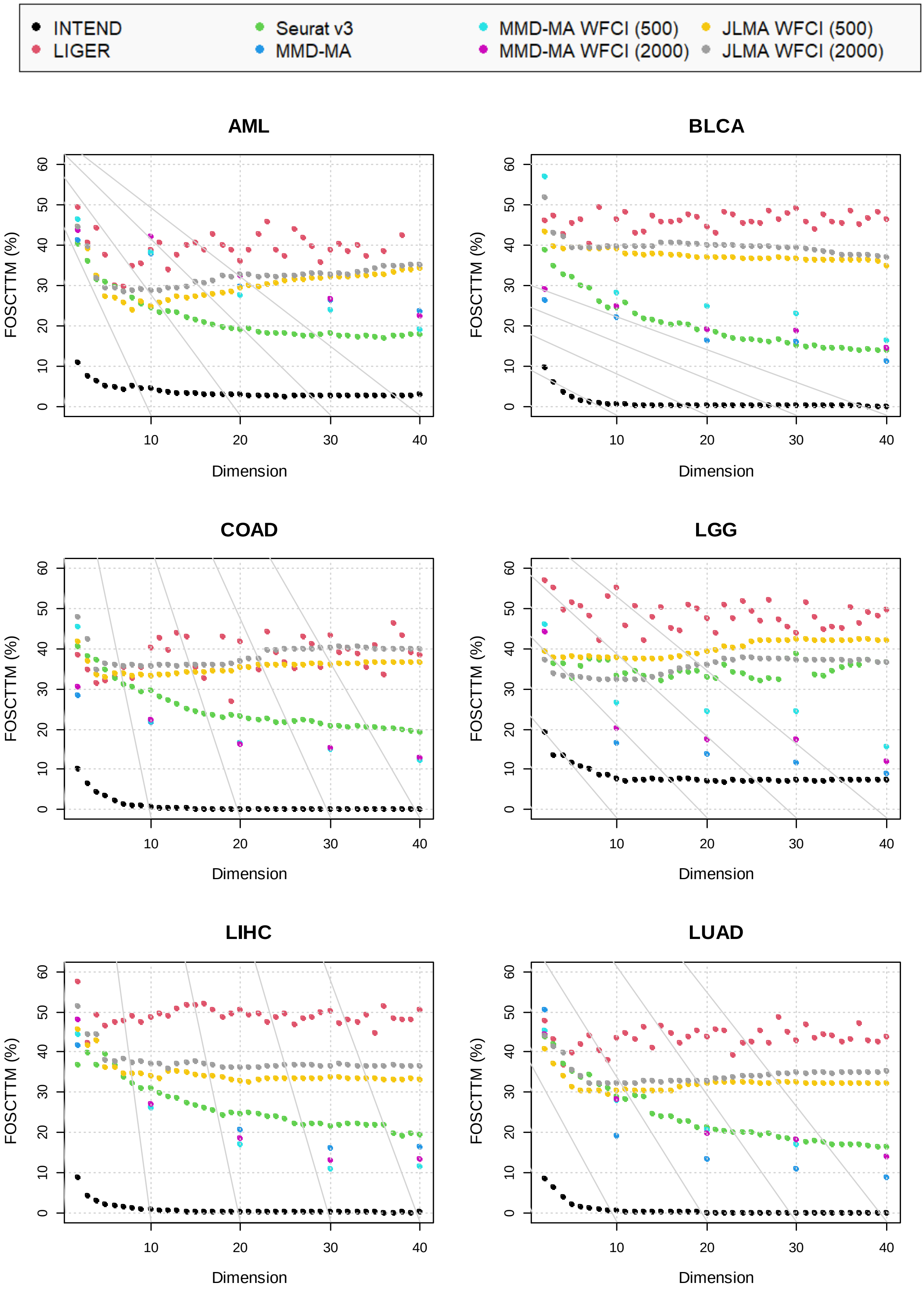
INTEND
LIGER
Seurat v3
MMD-MA
MMD-MA WFCI (500)
MMD-MA WFCI (2000)
JLMA WFCI (500)
JLMA WFCI (2000)
AML
BLCA
COAD
LGG
LIHC
LUAD
FOSCTTM (%)
Dimension
0
10
20
30
40
50
60

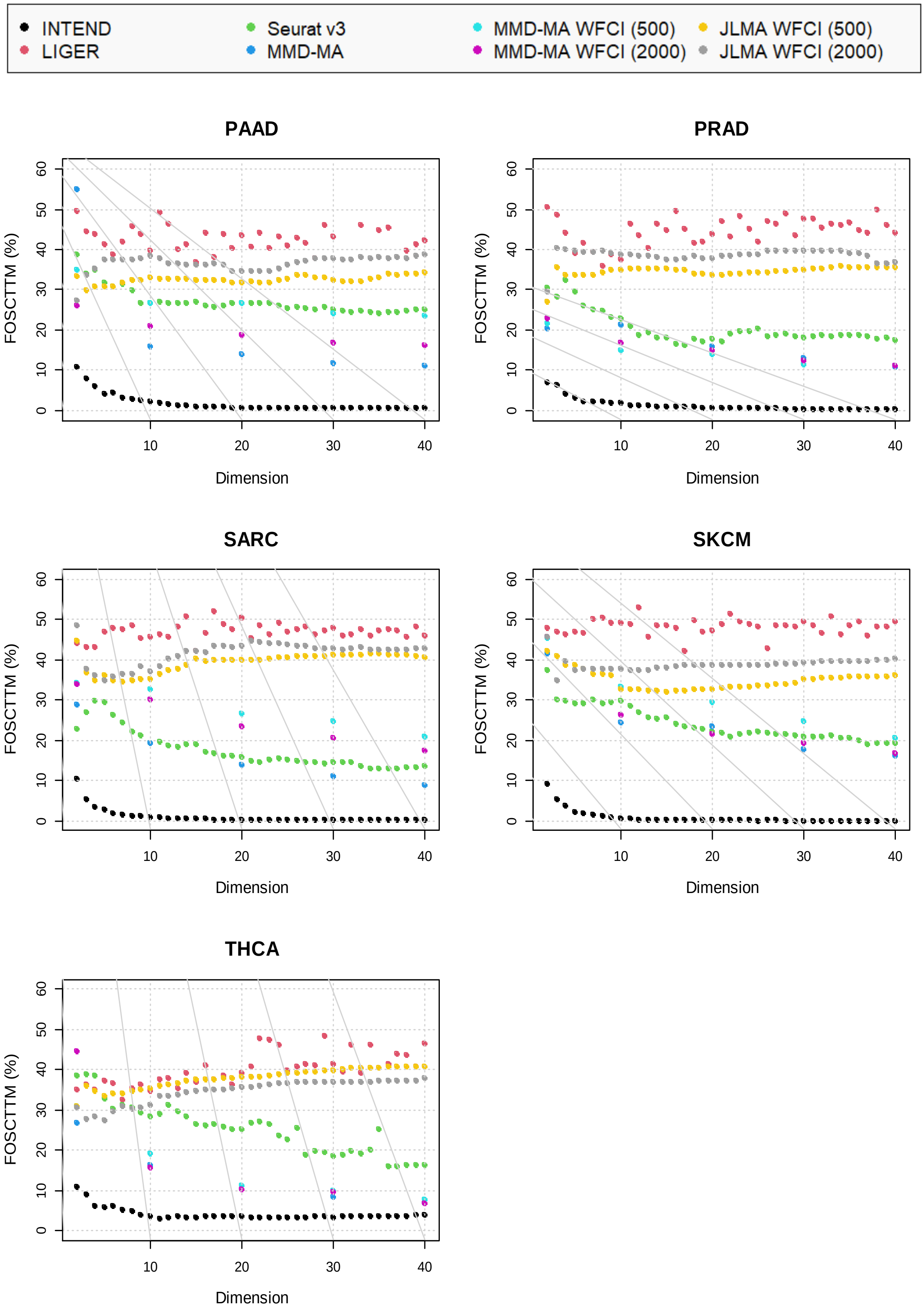


**Supplementary Figure 5: Performance of the algorithms as a function of the projected dimension on eleven TCGA cancer datasets.** Average FOSCTTM score versus the shared space dimension. The numbers 500 and 2000 in parenthesis denote the number of selected genes in the WFCI runs of MMD-MA and JLMA. The results of MMD-MA include only $d = 2, 10, 20, 30, 40$, due to the long runtime of the algorithm.

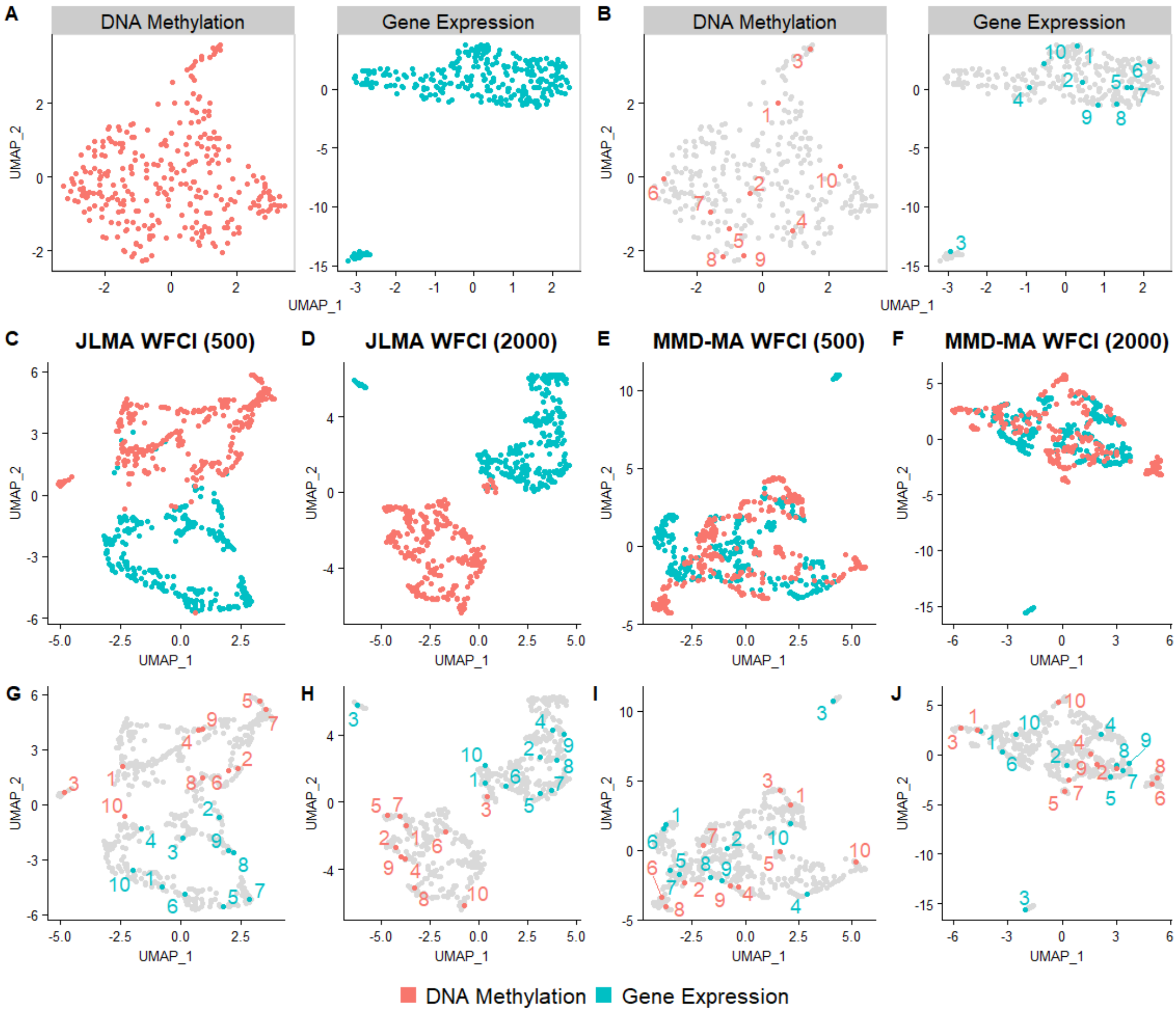


**Supplementary Figure 6.** The integration of gene expression and DNA methylation samples from the COAD dataset – results for JLMA and MMD-MA algorithms

(A) UMAP plots of the original data colored by omic.
(B) UMAP plots of the original data. To appreciate concordance between omics, ten samples were randomly chosen, and their matching points in both omics are labeled and colored by omic.
(C-J) UMAP plots of the samples after they were projected to a shared space by each algorithm, with a set of selected genes of size 500 and 2000. The samples are colored by omic (C-F) and the projection of the points from (B) are labeled in (G-J).

**Supplementary Figures 7-16.** Results of integration of GE and DM samples from all TCGA datasets listed in **Table 1**, excluding COAD. (A) UMAP plots of the original data. (B) The same plots as in A. To appreciate concordance between omics, ten samples were randomly selected, and their matching points in both omics were labeled. (C-F) UMAP plots of the samples after they were projected to a shared space by each algorithm. (G-J) The same plots as in C-F with the selected points labeled. In all plots colors correspond to omics.

**AML**

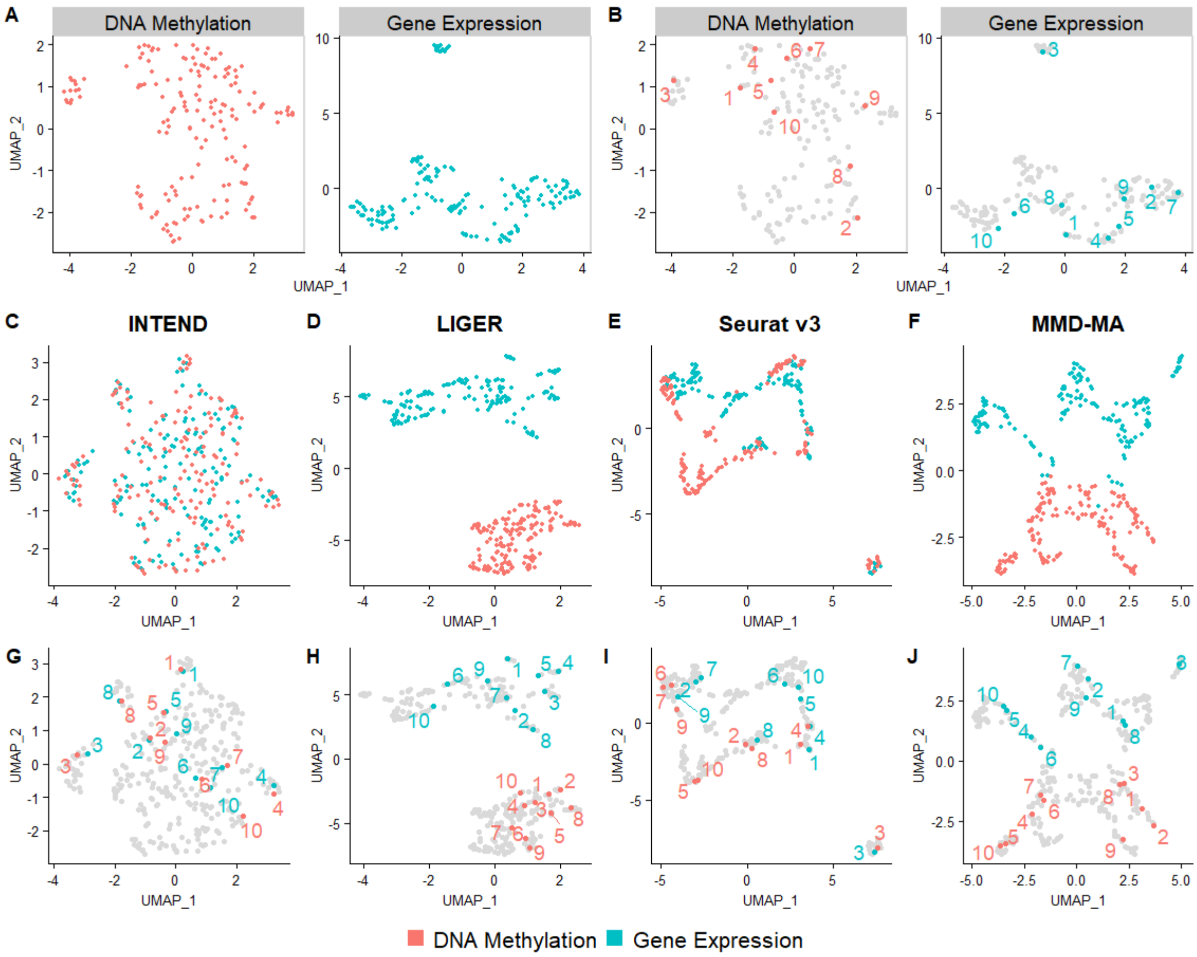

**BLCA**

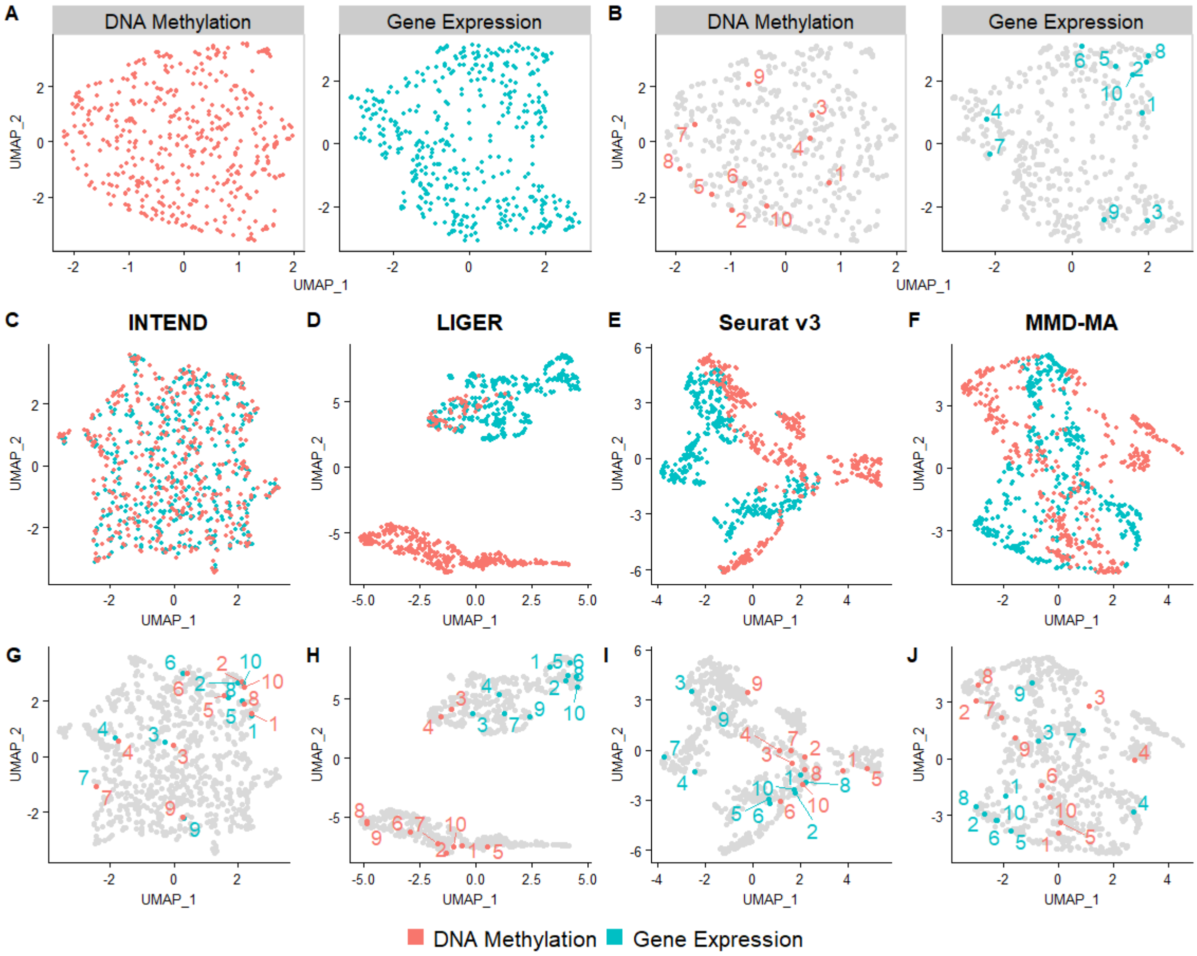
A
DNA Methylation
Gene Expression
B
DNA Methylation
Gene Expression
UMAP_1
UMAP_2
C
INTEND
D
LIGER
E
Seurat v3
F
MMD-MA
G
H
I
J
DNA Methylation
Gene Expression

**LGG**

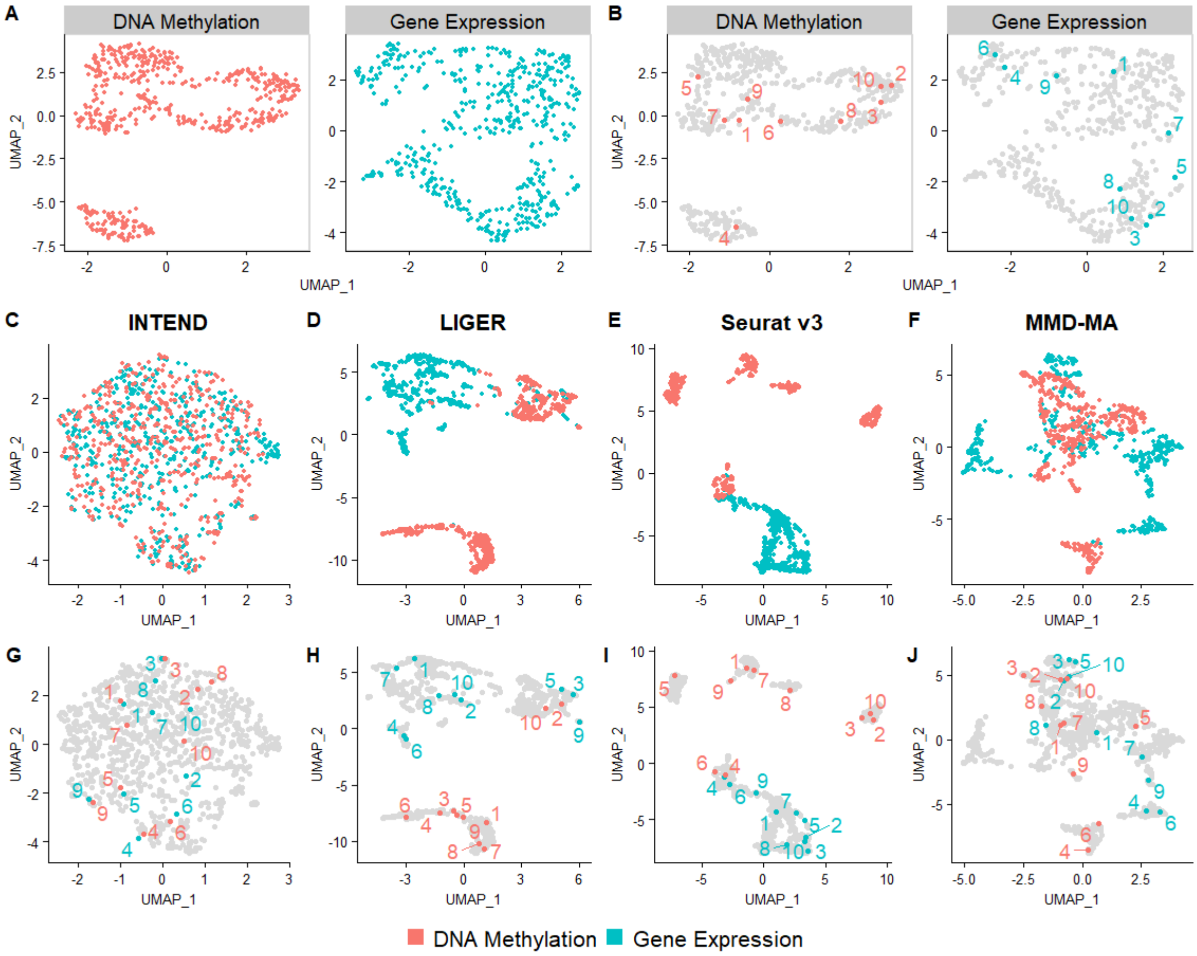

A
DNA Methylation
Gene Expression
B
DNA Methylation
Gene Expression
UMAP_1
UMAP_2
C
INTEND
D
LIGER
E
Seurat v3
F
MMD-MA
G
H
I
J
DNA Methylation
Gene Expression

**LIHC**

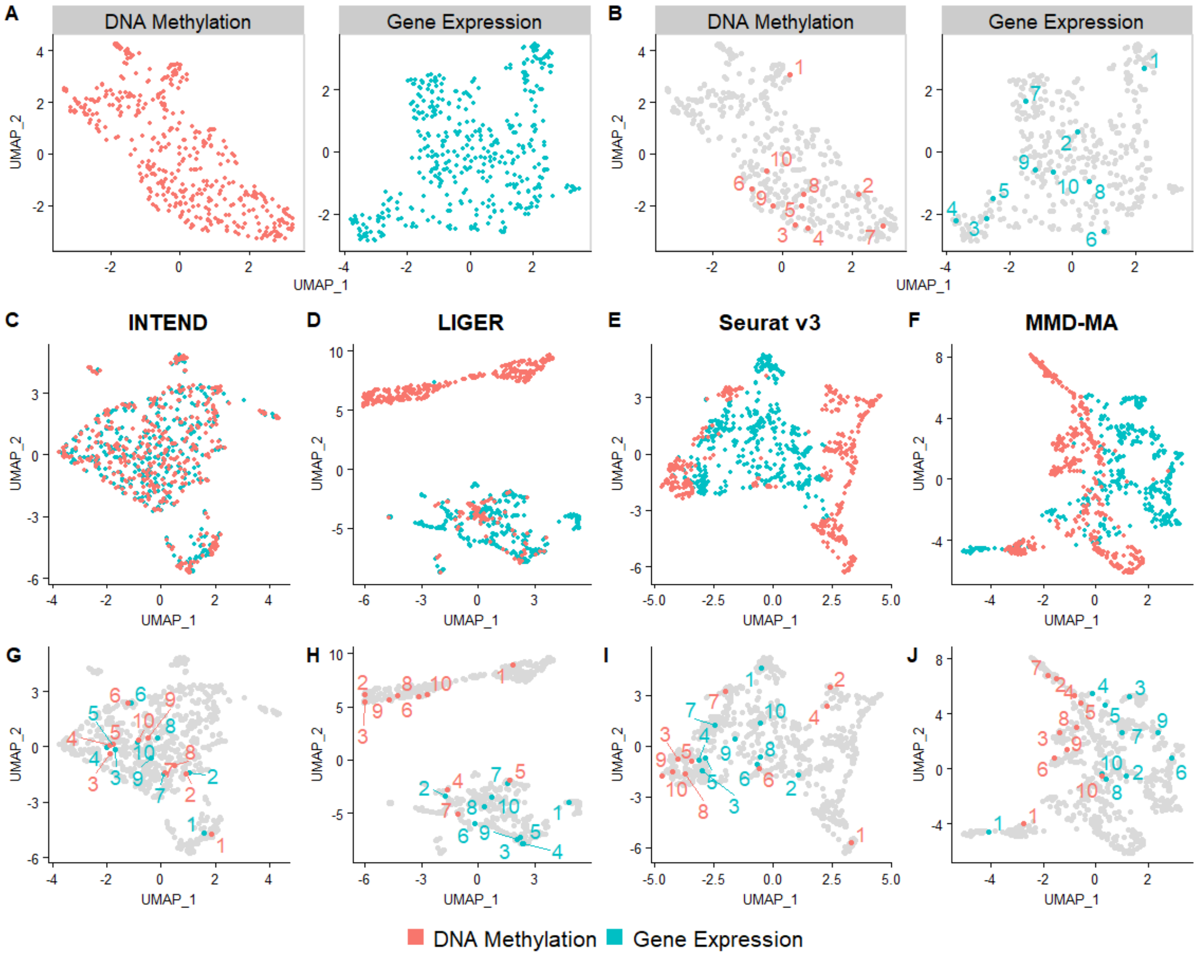

A
DNA Methylation
Gene Expression
B
DNA Methylation
Gene Expression
UMAP_1
UMAP_2
C
INTEND
D
LIGER
E
Seurat v3
F
MMD-MA
G
H
I
J
DNA Methylation
Gene Expression

**LUAD**

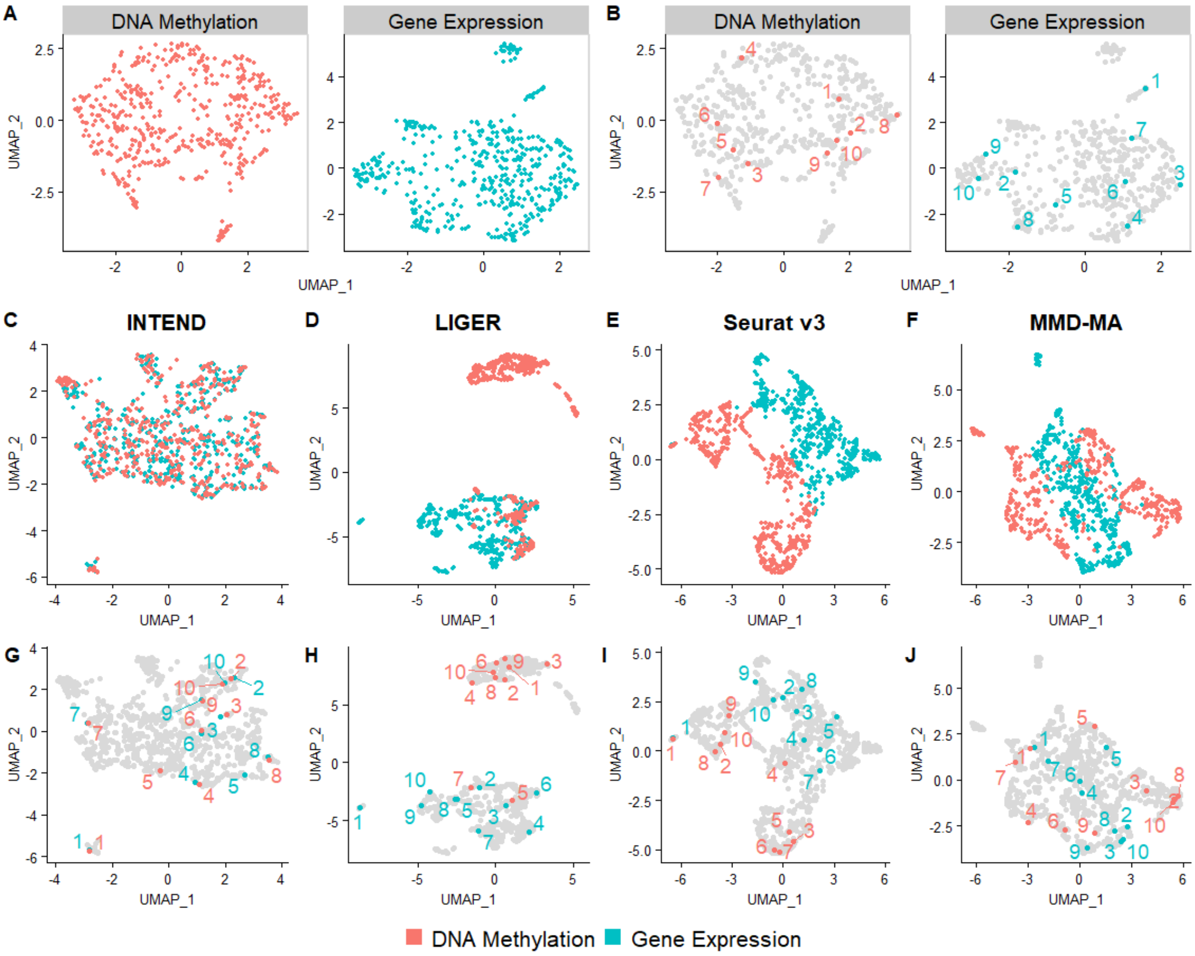
A
DNA Methylation
Gene Expression
B
DNA Methylation
Gene Expression
UMAP_1
UMAP_2
C
INTEND
D
LIGER
E
Seurat v3
F
MMD-MA
G
H
I
J
DNA Methylation
Gene Expression

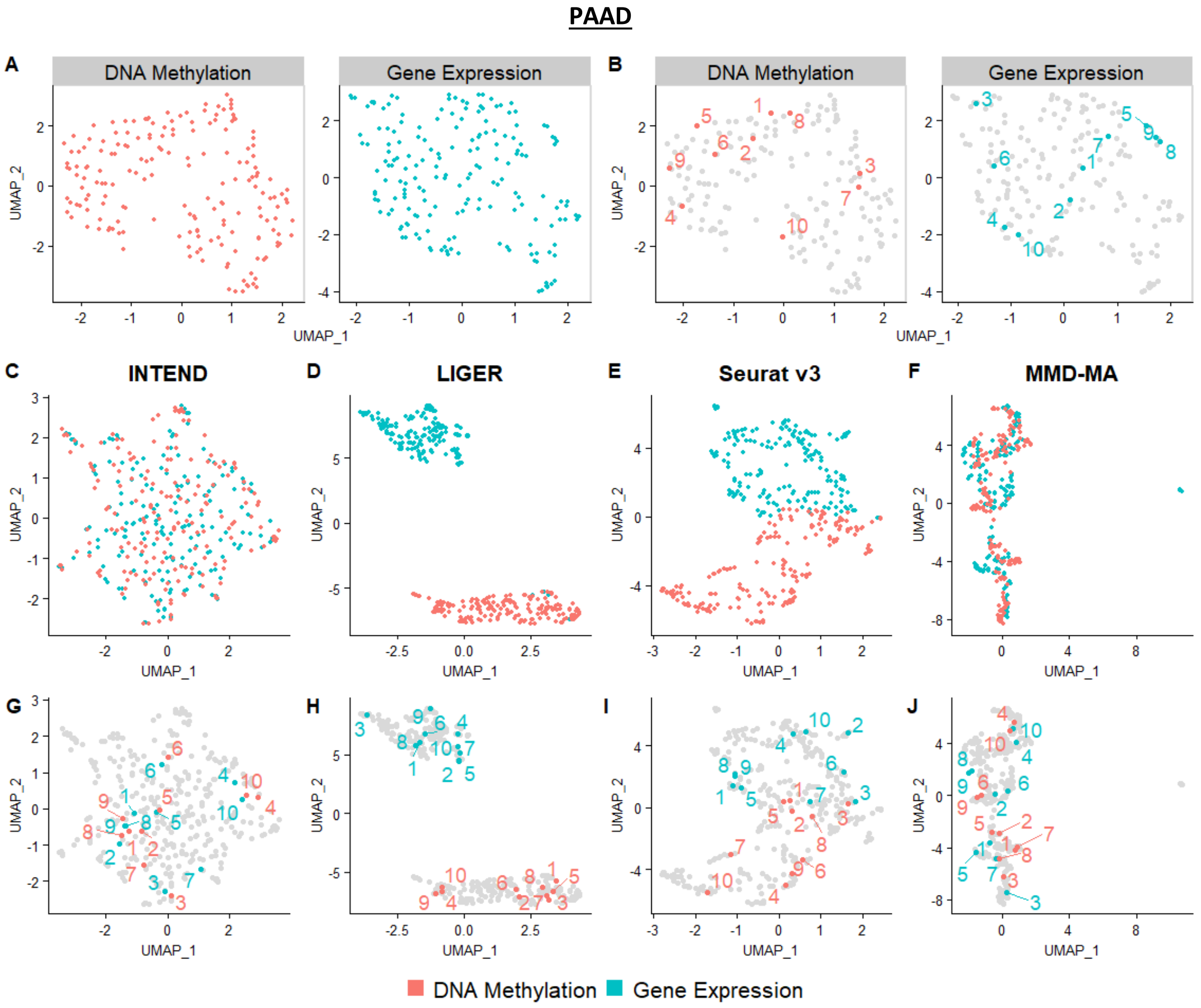

PAAD
A
DNA Methylation
Gene Expression
B
DNA Methylation
Gene Expression
UMAP_1
UMAP_2
C
INTEND
D
LIGER
E
Seurat v3
F
MMD-MA
G
H
I
J
DNA Methylation
Gene Expression

**PRAD**

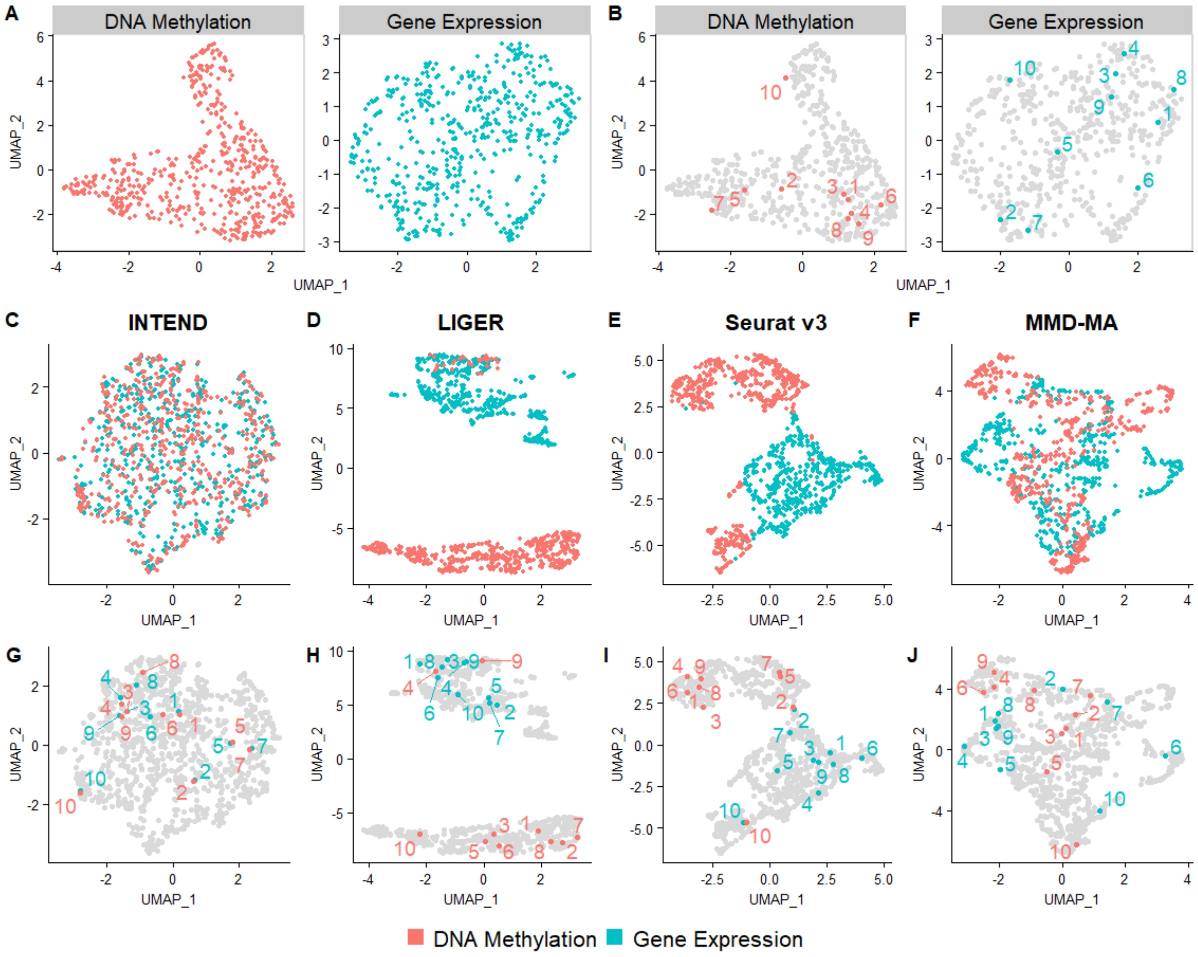
A
DNA Methylation
Gene Expression
B
DNA Methylation
Gene Expression
C
INTEND
D
LIGER
E
Seurat v3
F
MMD-MA
G
H
I
J
UMAP_1
UMAP_2
DNA Methylation
Gene Expression

**SARC**

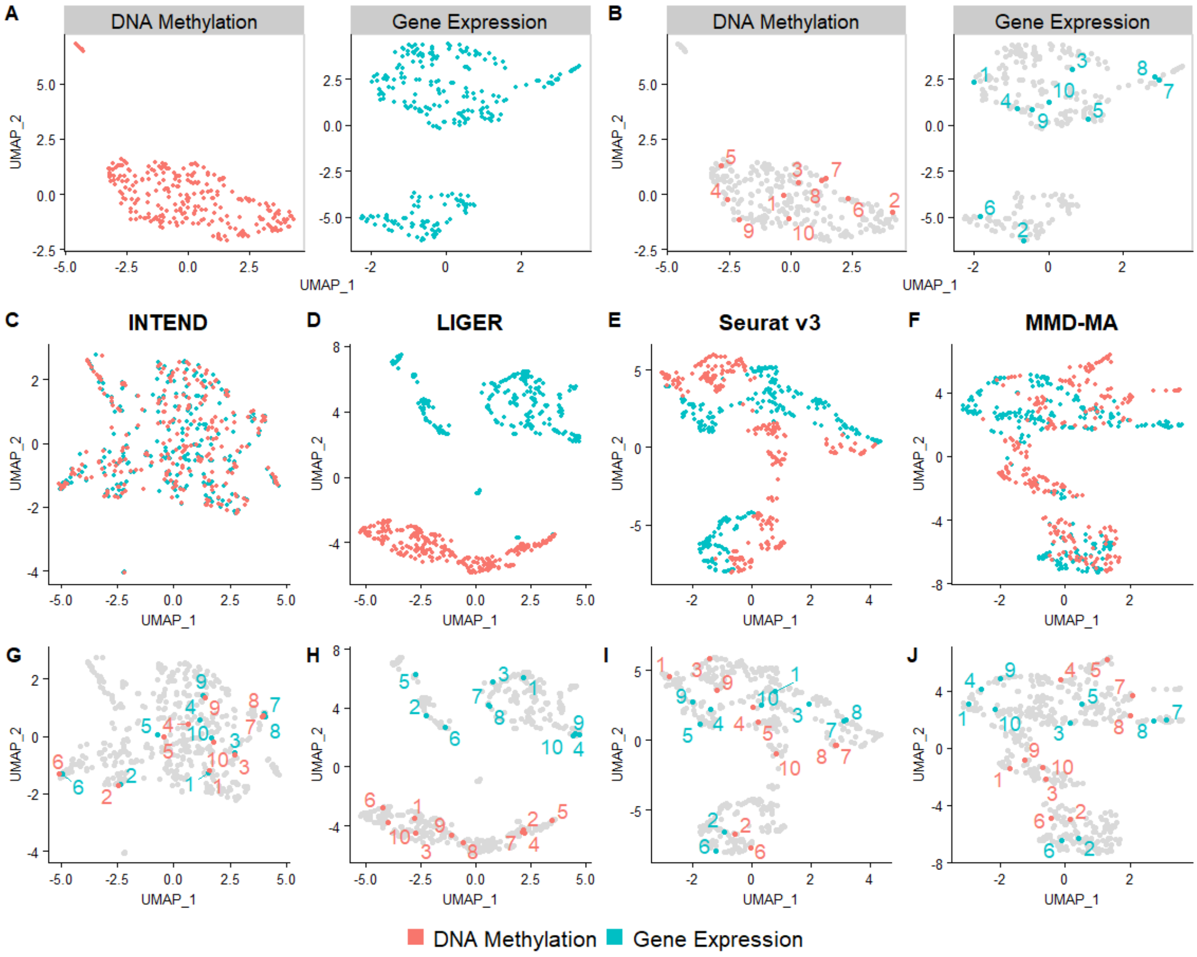

A
DNA Methylation
Gene Expression
B
DNA Methylation
Gene Expression
UMAP_1
UMAP_2
C
INTEND
D
LIGER
E
Seurat v3
F
MMD-MA
G
H
I
J
DNA Methylation
Gene Expression

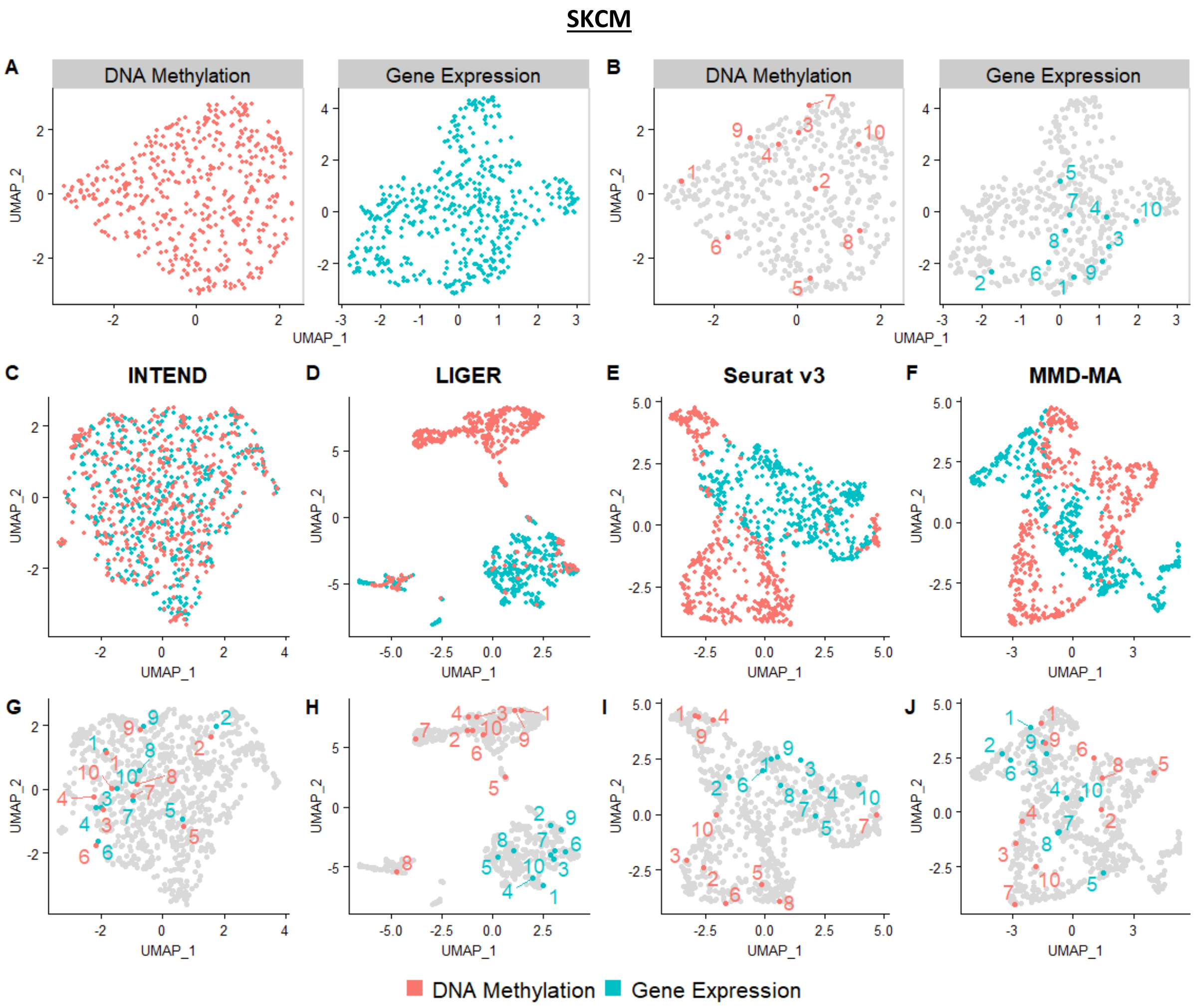

SKCM
A
DNA Methylation
Gene Expression
B
DNA Methylation
Gene Expression
C
INTEND
D
LIGER
E
Seurat v3
F
MMD-MA
G
H
I
J
UMAP_1
UMAP_2
DNA Methylation
Gene Expression

**THCA**

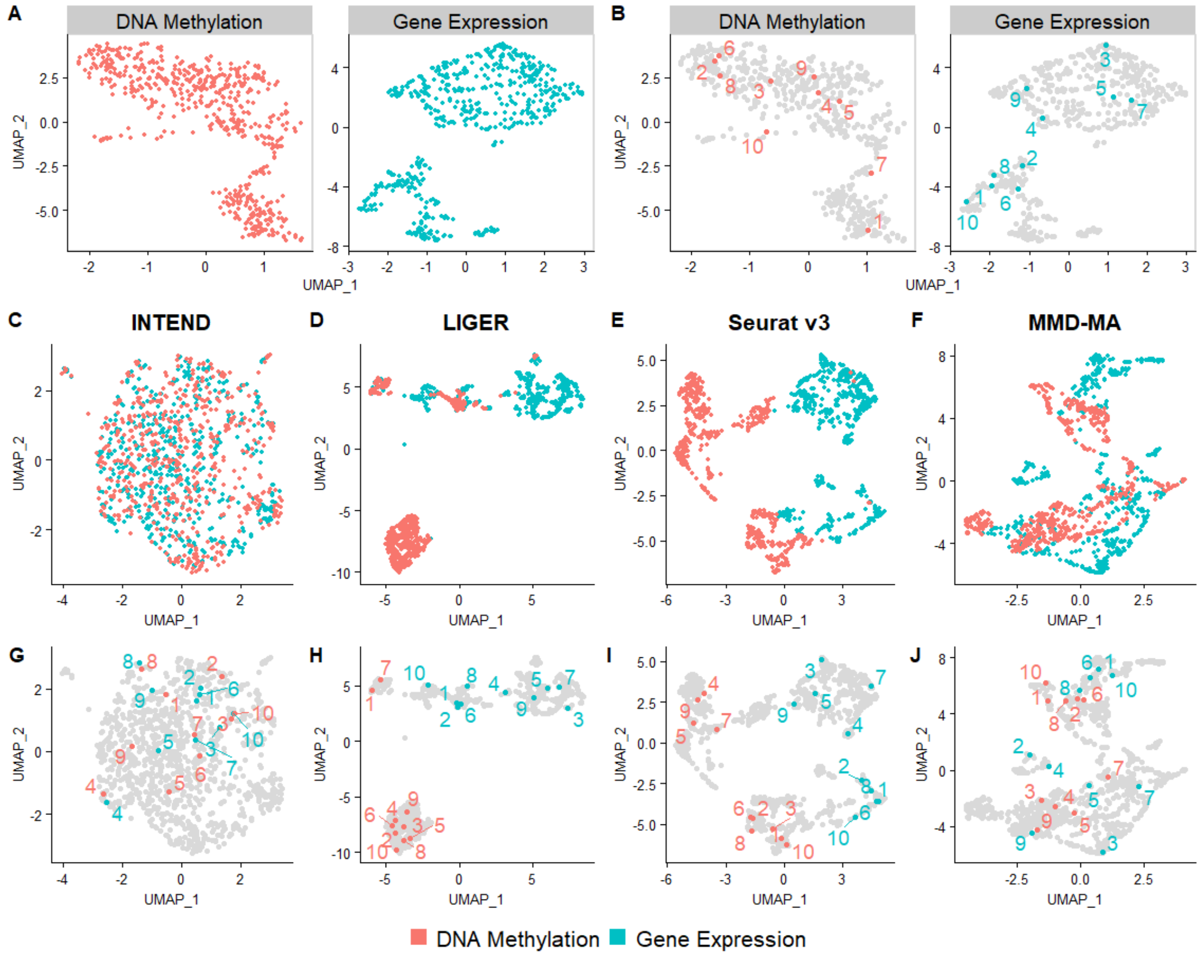
A
DNA Methylation
Gene Expression
B
DNA Methylation
Gene Expression
UMAP_1
UMAP_2
C
INTEND
D
LIGER
E
Seurat v3
F
MMD-MA
G
H
I
J
DNA Methylation
Gene Expression

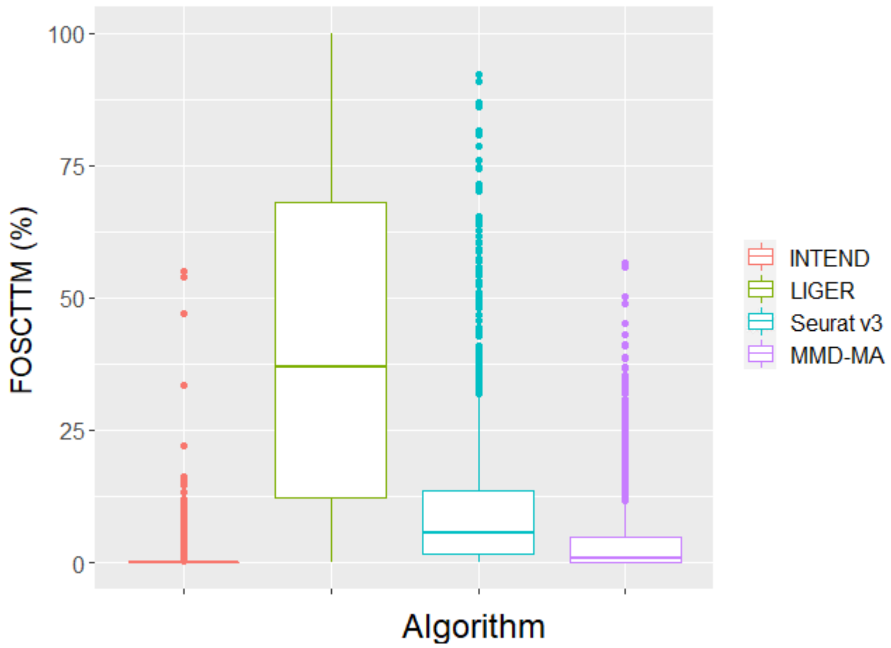


**Supplementary Figure 17.** FOSCTTM scores of the integration of four cancer datasets: COAD, LIHC, SARC and SKCM, simultaneously, by INTEND, LIGER, Seurat v3, and MMD-MA.

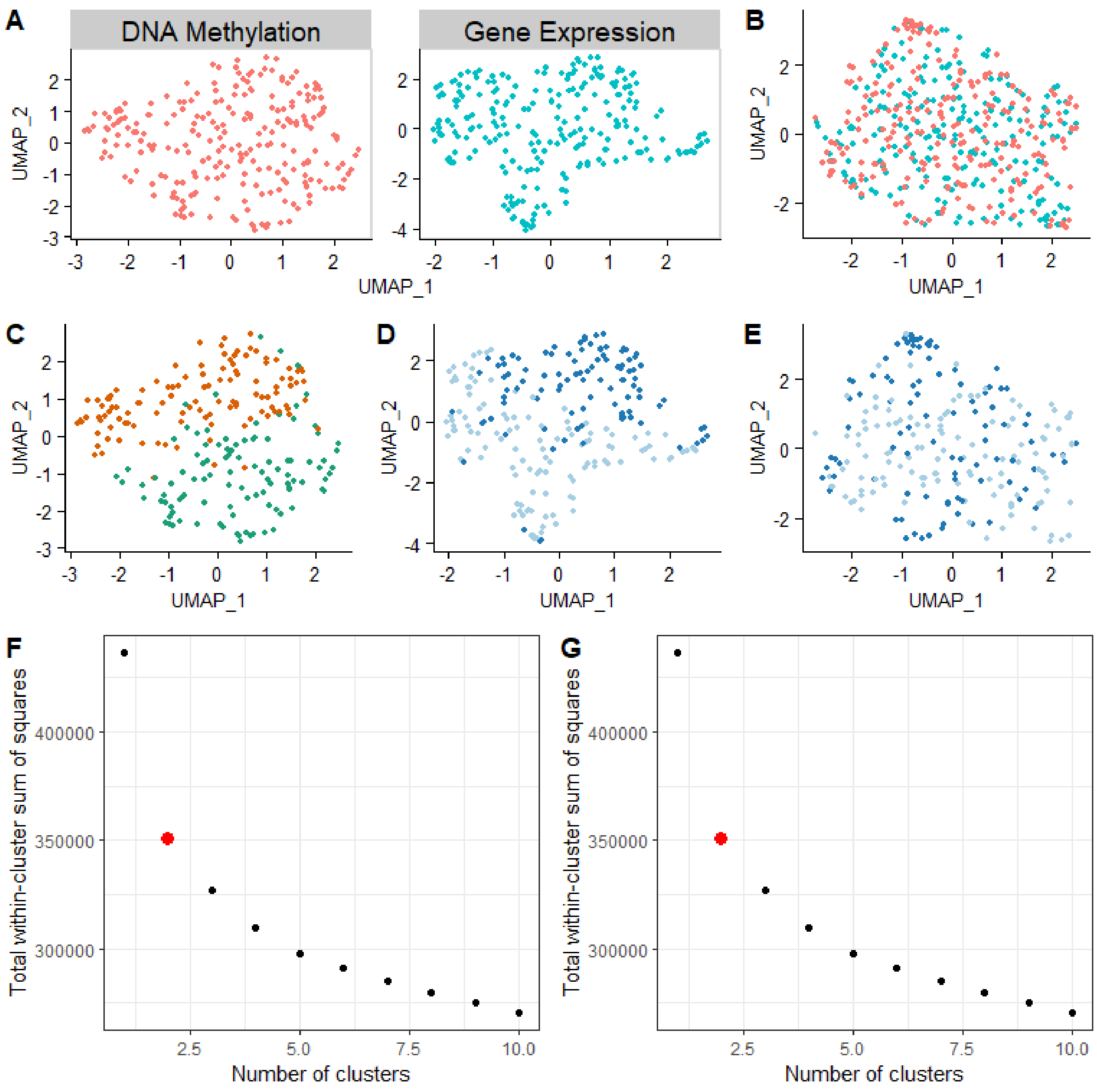


**Supplementary Figure 18.** SKCM clustering

(A) UMAP plots of each original omic data.
(B) UMAP plot of INTEND sample projections into the shared space colored by omic
(C) k-means clustering of the original DM samples with $k = 2$, shown on the same plot as in (A). Samples are colored according to their clusters.
(D) k-means clustering of the original GE samples with $k = 2$, shown on the same plot as in (A). Samples are colored according to their clusters.
(E) Integration-based clustering of the methylation sample embeddings into the shared space (EP, the pink points in (B)), shown on the same plot as in (B). Samples are colored according to their assigned cluster from (D). Each sample was assigned by majority voting to the cluster most represented among the five GE embeddings closest to its matching EP representation in the shared space.
(F) The total within-cluster sum of squares versus the number of clusters, for clustering DM data.
(G) The total within-cluster sum of squares versus the number of clusters, for clustering GE data.
The points with the maximum curvature are highlighted in red.
